# Inferring modularity in genotype-phenotype maps using patterns of epistasis and pleiotropy

**DOI:** 10.64898/2026.11.03.764076

**Authors:** Nicholas Lawson, Grace Brophy, Caroline M. Holmes, Anna Seigal, Michael M. Desai, Samantha Petti

**Affiliations:** Department of Physics, Harvard University, Cambridge, MA 02138; Department of Mathematics, Tufts University, Medford, MA 02155; Department of Organismic and Evolutionary Biology, Harvard University, Cambridge, MA 02138; John A. Paulson School of Engineering and Applied Sciences, Harvard University, Cambridge, MA 02138; Department of Computer Science, Tufts University, Medford, MA 02155

## Abstract

Methods for inferring the relationship between genotype and phenotype often search for low-dimensional structure, motivated by the hypothesis that genetic loci can be organized into a small number of groups representing biological processes. These methods typically group genetic loci based on which other loci they interact with (epistasis) or based on how their effects covary across environmental conditions (patterns of pleiotropy). Here, we present two complementary methods that combine epistasis and pleiotropy to infer modular structure. The first, *Sparse, Low-dimensional, Epistatic Decomposition* (SLED), parametrizes the genotype-phenotype map in terms of a small number of modules: loci contribute sparse additive and epistatic effects to modules, and modules contribute sparse additive and epistatic effects to multiple phenotypes. The second method, *Graph Tensor Decomposition* (GTD), represents the epistatic interactions in each environment as a combination of shared basis graphs, each capturing a pattern of epistasis that recurs across environments. Since GTD does not use additive effects, learned graphs are not restricted to include only loci whose additive effects covary across environments. We validate SLED and GTD on synthetic data, then apply both methods to an experimental dataset consisting of the growth rates of yeast segregants from an F1 cross, measured across 101 environments with a hub-and-spoke structure. We evaluate the biological structure of the inferred modules and show that both models generalize well to held-out environments. Finally, we compare the modules inferred by SLED, GTD, and a previously established method that infers sparse modular structure from a locus-by-environment matrix of additive effects (SSD) and find that all three methods yield highly similar modular architectures. This suggests that the genetic architecture underlying this dataset is consistent across additive effects (SSD), epistatic effects (GTD), and both jointly (SLED). Together, our methods provide a framework for combining two sources of information, epistasis and pleiotropy, to infer interpretable low-dimensional structure in genotype-phenotype maps, expanding on previous work that has traditionally treated each independently.

---

The structure of genetic interactions has long been used as the basis for inferring patterns of biological organization. A guiding hypothesis underlying many of these methods is that genotype-phenotype (GP) maps are composed of low-dimensional “modules,” which represent different biological processes and pathways (1). One group of approaches uses epistatic interactions, mapped extensively in model organisms (2–7), to identify these functional modules. Specifically, numerous studies use clustering methods to form groups of genes that either interact strongly (8), or have similar interaction profiles across all other genes (9). Comparisons across a small number of conditions reveal that these epistatic patterns vary systematically with environment (10–12).

Another class of approaches for identifying modularity focuses on pleiotropy (13–15), which we define here as the influence of a single genetic locus on a given phenotype across different environments. Modern high-throughput screens have shown that pleiotropy is widespread in both humans and model organisms (6, 16–19). One way to infer modular structure using pleiotropy is to apply matrix decomposition methods to the *G* × *E* additive effects matrix (20–22). Many such approaches (e.g. Sparse Structure Discovery (SSD) (23) and Webster (21)) build on the classic matrix decomposition technique SVD (16), adding regularization penalty terms that reflect biological assumptions. These additive methods are often interpretable, but neglect epistasis by construction. In contrast, nonlinear machine learning approaches can sometimes yield greater predictive accuracy, but often at the expense of biological interpretability (24, 25).

Here, we introduce two complementary methods for inferring modular structure in GP maps that leverage patterns of both epistasis and pleiotropy (Fig. 1). The first method, *Sparse, Low-dimensional, Epistatic Decomposition* (SLED), identifies a low-dimensional, nonlinear decomposition of the GP map in which loci contribute sparse additive and epistatic effects to a small number of intermediate modules. Phenotypes in different environments are then jointly modeled as sparse linear and pairwise combinations of these modules. SLED is a nonlinear generalization of the *G* × *E* matrix decomposition method SSD, allowing epistasis within and between modules, which we refer to as intraand inter-modular epistasis, respectively.

**Fig. 1.**
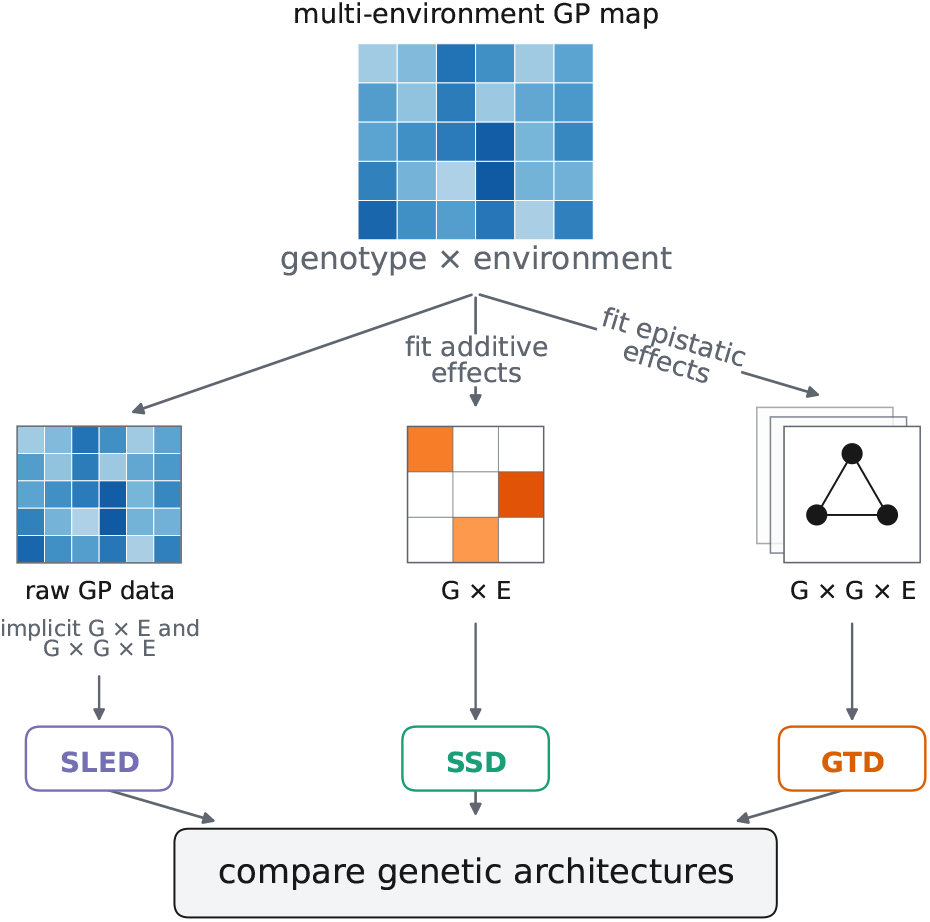
SLED and GTD incorporate the effects of epistasis and pleiotropy to infer modularity in multi-environment genotype-phenotype (GP) maps. SLED directly analyzes GP data. The linear method SSD (23) inputs a pre-fit *G* × *E* additive effects matrix, and GTD inputs a pre-fit *G* × *G* × *E* tensor of epistatic effects across environments. We then compare the genetic architectures inferred by the three models.

SLED allows interactions between loci in a module only when they both contribute additive effects to that module, which prohibits interactions between loci with decoupled additive effects across environments. To address this, our second method, *Graph Tensor Decomposition* (GTD), identifies shared patterns of pairwise epistasis across phenotypes without considering additive effects. It decomposes a tensor of *G* × *G* × *E* interactions such that these interactions for each phenotype are expressed as a sparse linear combination of a small set of shared basis graphs, each of which represents a pattern of interactions between the loci. Our methods belong to a broader class of approaches that model structure in GP maps, which we return to in the Discussion.

These complementary methods let us ask how the inferred modular structure depends on the choice of an additive, epistatic, or joint model of genetic architecture. We explore this by comparing the modules inferred by SSD, GTD, and SLED on an experimental dataset consisting of the growth rates of yeast segregants from an F1 cross, measured across many structured environments. We find that, despite the substantial differences between the three methods, the emergent genetic architectures are strikingly consistent, providing support for the modularity hypothesis.

The structure of the paper is as follows. In Sections 1 and 2, we present each of the models and validate them on synthetic data. In Section 3, we apply both models to the empirical GP map described above. We interpret the biological organization learned by each model and show that the inferred genetic architectures successfully predict growth rates in novel environments. Finally, by comparing SSD, GTD, and SLED, we ask how the inferred modules depend on each method’s input: additive effects, epistatic interactions, or both.

## 1. SLED: Sparse, Low-dimensional, Epistatic Decomposition

### 1.1. Description of SLED

*Sparse, Low-dimensional, Epistatic Decomposition* (SLED) is a model that predicts phenotypes from genotypes (Fig. 2). SLED jointly predicts *E* phenotypes for each individual, which could be *E* different traits or, as in this paper, growth rates in *E* environments. We assume that there exists low-dimensional structure in the GP map, such that it can be described by a small number of intermediate modules or core processes. We also assume that the maps from loci to modules and modules to environments are sparse.

**Fig. 2.**
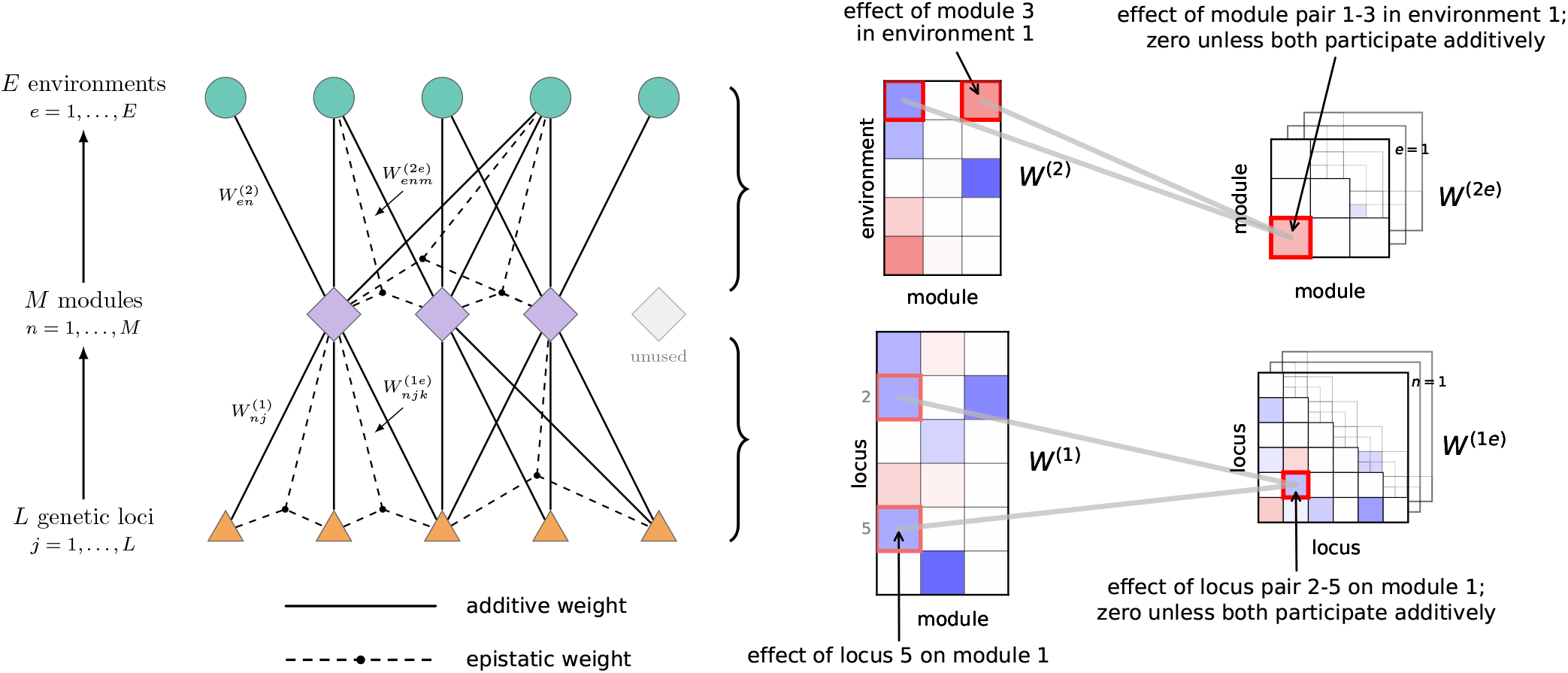
Overview of SLED. Left: SLED is a sparse, low-dimensional map from genotype to phenotype that incorporates epistasis. Loci (orange triangles) contribute additive and epistatic effects to a small number of shared modules (purple diamonds); the modules contribute additive and epistatic effects to the phenotype in each environment (teal circles). Right: the learned weights are sparse and obey a “linked epistasis” constraint: epistatic interactions are allowed only between pairs of loci (modules) contributing additive effects to a given module (environment).

Intuitively, each locus affects only a small number of modules, and, in any given environment, an organism’s phenotype is determined by a subset of the modules.

Previous work proposed a linear method, *Sparse Structure Discovery* (SSD), for obtaining a sparse, low-rank decomposition of a GP map (23). SSD infers two sparse maps: the first from loci to a small number of modules, and the second from those modules to fitness effects across environments; this constitutes a low-rank approximation to the full matrix of fitness effects, where the modules are inferred from pleiotropic information. SSD also allows for the number of modules and the level of sparsity at each layer to be selected interpretably from the data.

Here, we extend SSD in two ways. First, SLED operates directly on GP data, rather than requiring a pre-inferred matrix of fitness effects (though SLED can also operate directly on tensors of fitness effects; see SI S1.2). Second, SLED generalizes SSD to allow for epistasis, both between loci within a module (intra-modular epistasis) and between modules (inter-modular epistasis). We assume that loci (modules) can interact in a module (environment) only if they both participate additively in that module (environment). In other words, we allow interactions within and between modules, but not otherwise—a “linked epistasis” constraint. Following SSD, we assume that both additive and epistatic effects are, to varying degrees, sparse.

Concretely, for a genotype with biallelic loci *gj* ∈ {−1, +1}, *j* = 1, …, *L*, and phenotypic predictions 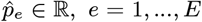, *e* = 1, …, *E*, we write

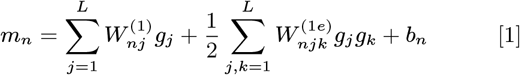

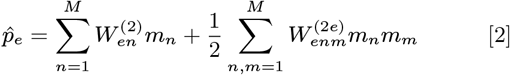

such that each module *mn, n* = 1, …, *M* is a linear plus pairwise function of the loci, and each predicted phenotype is a linear plus pairwise function of the modules. Four hyperparameters {*λ*1, *λ*1*e, λ*2, *λ*2*e*}, one for each of the weight tensors

{*W* ^(1)^, *W* ^(1*e*)^, *W* ^(2)^, *W* ^(2*e*)^}, encourage sparsity in the weights (see SI S1.1 for the loss function and optimization details). The loss function also enforces the linked epistasis assumption.

Various extremes of the four-dimensional hyperparameter space correspond to previously established methods. Strong epistatic but weak linear regularization effectively reduces SLED to SVD, the matrix decomposition method with the lowest reconstruction error; strong epistatic and moderate linear regularization reduces it to SSD. At a fixed, moderate level of linear regularization, the model allows for two distinct notions of epistasis. When only intra-modular epistasis (*W* ^(1*e*)^) is strongly regularized, all loci in a module share a common interaction pattern with loci in another module, analogous to clustering genes that have similar effects on a phenotype (16, 23). On the other hand, when only inter-modular epistasis (*W* ^(2*e*)^) is strongly regularized, a module is a set of loci that interact with one another, analogous to clustering genes based on their gene-gene interaction profiles (8). By examining models that achieve the same predictive accuracy but trade off sparsity in different regimes, we can obtain a family of models that emphasize different kinds of interactions (see SI S1.1.5).

### 1.2. Validating SLED on synthetic data

We first validated SLED on synthetic phenotypes drawn from different generative processes designed to isolate the effects of sparsity and modularity (Fig. 3). These weights were 1) sparse and modular, 2) dense and modular, and 3) sparse and non-modular. In the sparse, modular case, the weights were sparse, the number of modules was small, and the existence probability of an epistatic weight depended on the product of the corresponding additive weights, matching the linked epistasis constraint of SLED. To remove sparsity while retaining modularity, we used dense weights while keeping the number of modules small. To instead remove modularity while retaining sparsity, we set the number of modules equal to the number of loci, eliminating low-dimensionality in the GP map. We also generated 4) a set of phenotypes determined entirely by random fifth-order interactions, which cannot be modeled by SLED. In all cases, we added Gaussian noise of 10% of the magnitude of phenotypic variation to the synthetic phenotypes.

**Fig. 3.**
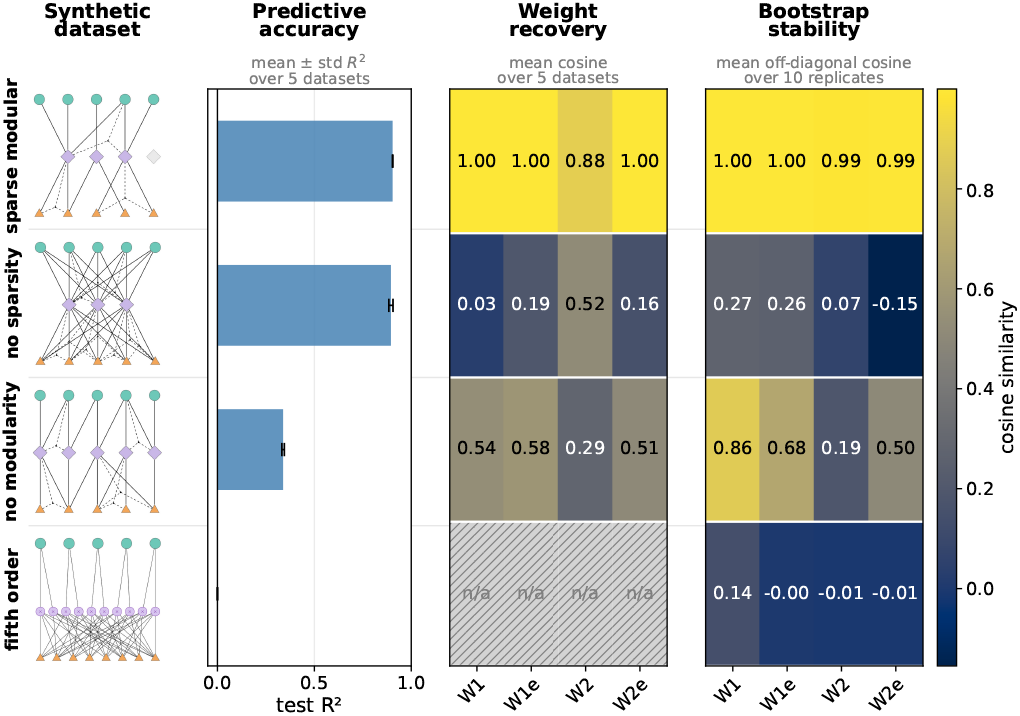
SLED applied to synthetic data. Rows correspond to different sets of synthetic GP data. All datasets have *L* = 100 loci and *E* = 20 phenotypes with 10% Gaussian noise added. First column: schematic illustrations of the generative processes, as described in the text. Second column: average *R*^2^ of trained models on test phenotypes; error bars over five simulated datasets. Third column: for each set of weights, mean cosine between the learned and generative weights, averaged over five datasets. For the non-modular weights generated with 100 modules (third row), the cosine is computed relative to the 10 best-match modules. The fifth-order phenotypes (last row) were not generated with weights matching the SLED architecture, so no recovery statistics are computed. Fourth column: for each set of weights, mean cosine between replicate model fits on the same dataset with different random initializations, averaged over 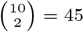 model pairs.

For each dataset, we tested the ability of SLED to recover the known generative weights, the predictive power of the model, and the stability of the models across replicate fits with different random seeds. We hypothesized that, if there existed sparse, modular structure in the data, the model would reliably find it, thus leading to consistent models across runs. By contrast, if the generative process did not match the structure that SLED assumes, we hypothesized that the model would return more random, and thus less similar, solutions across training runs.

SLED achieved excellent recovery, predictive accuracy, and stability on the sparse, modular dataset. The model was able to predict the dense, modular dataset well, but struggled to recover the ground truth weights. This occurred because, over replicate training runs, the model found many different non-sparse solutions that explained the data well, none of which entirely matched the ground truth. The model could neither predict nor recover the sparse, non-modular dataset well because it was forced to compress a high-dimensional GP map into a low-dimensional representation. Finally, as expected, the model was completely unable to predict the fifth-order phenotypes. These results confirmed that SLED can reliably detect sparse, low-dimensional, nonlinear structure in GP maps and, importantly, that SLED does not invent such structure in datasets that do not exhibit it.

## 2. GTD: Graph Tensor Decomposition

### 2.1. Description of GTD

Our *Graph Tensor Decomposition* (GTD) is a model that finds a low-rank decomposition of epistatic interactions across environments (Fig. 4). Unlike SLED, GTD does not account for linear effects; instead, this method asks if a modular genetic architecture is discoverable from epistatic interactions alone. As such, GTD does not directly operate on GP data, but rather on pre-inferred epistatic weights that capture *G* × *G* × *E* interactions.

**Fig. 4.**
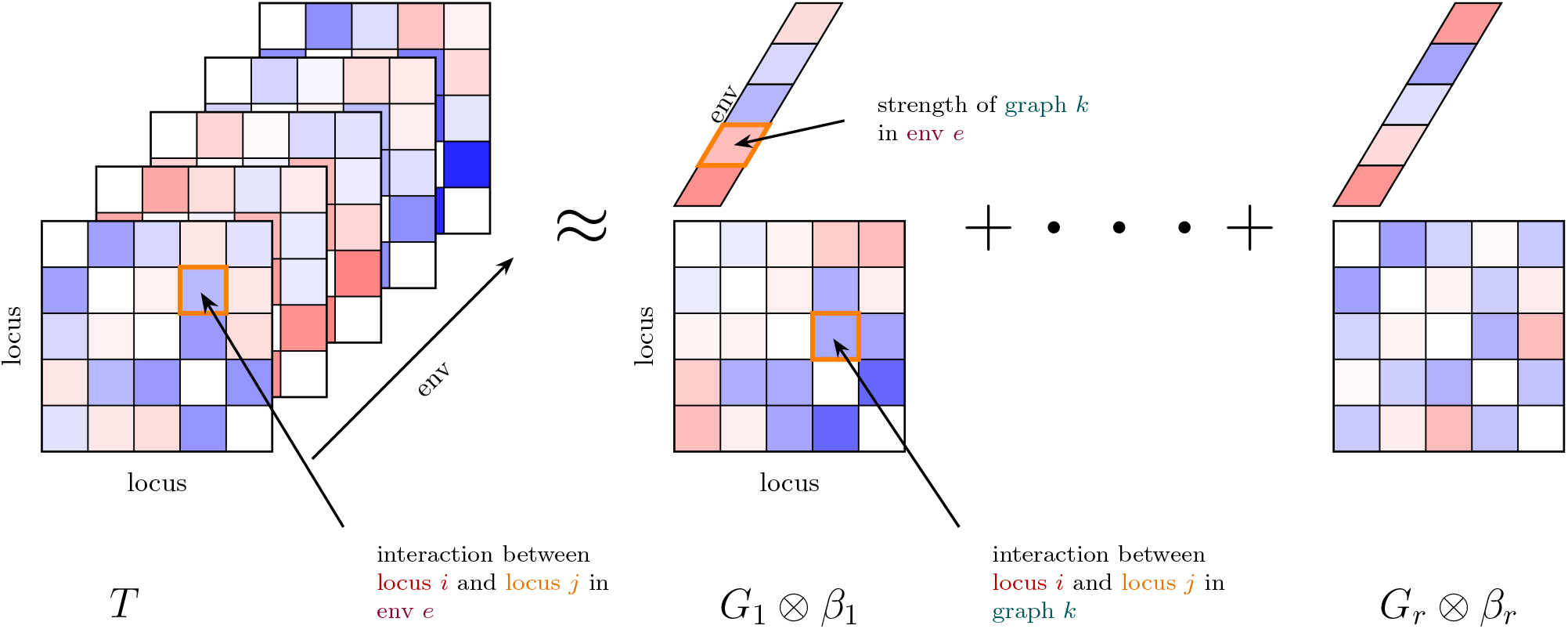
Overview of GTD. GTD decomposes a tensor of epistatic interactions *T* into a small set of sparse basis graphs 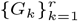, each with sparse activation across environments. Locus-by-locus matrices represent the learned graphs, and the accompanying length-*E* vectors represent their per-environment weights 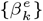; ⊗ denotes the outer product between graphs and weight vectors.

Concretely, the input to GTD is an *L* × *L* × *E* tensor *T* of epistatic interactions across environments. Our goal is to decompose *T* into a set of *r* shared basis graphs 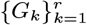 (i.e. *L* × *L* matrices of pairwise interactions) and per-environment weights 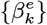 such that 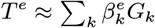 for each environment *e*. GTD belongs to a broader family of tensor decomposition algorithms (26–30). However, usual tensor decomposition imposes too narrow a structure, enforcing rank-one components. For example, MCPCA (31) finds a strength vector *v* such that *G*_*k*_ = *vv*^*T*^, i.e. that interactions between loci are products of per-locus weights. GTD instead allows *G*_*k*_ to be any sparse graph, assuming that a small number of loci participate in each graph, and that these graphs are active across a subset of environments. We compare GTD to MCPCA in SI S2.8.

Formally, we derive our loss function from a Bayesian perspective. We impose sparsity-inducing priors on graphs and weights. Following an Erdős–Rényi prior, loci participate with probability *p*, and interactions exist between loci with probability *q* in each graph *G*_*k*_. Most per-environment weights are expected to be zero, as graphs are active across only a small subset of all environments. Minimizing the following loss function

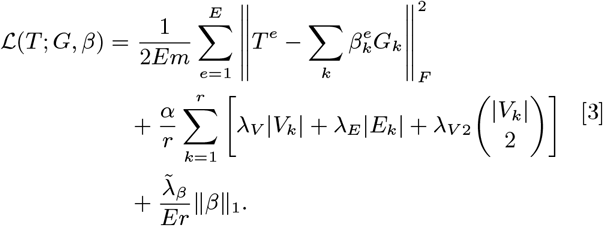

yields the maximum *a posteriori* (MAP) estimate of our Bayesian model, where 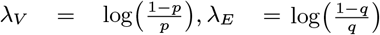, and *λV* 2 = − log(1 − *q*) for the graph vertex and edge probabilities *p* and *q* described above. Briefly, the first term in the loss is the mean-squared reconstruction error of the epistasis tensor, averaged over the *Em* unique pairwise entries, with 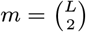; *α* sets the strength of the graph sparsity regularization on vertex sets *V*_*k*_ and edge sets *E*_*k*_ of each graph *G*_*k*_; and 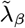 controls the sparsity of the per-environment weights (see SI S2.1 for more details).

### 2.2. Validating GTD on synthetic data

We validated GTD on synthetic data by constructing tensors from various underlying graph structures. We computed GTD’s ability to recover the underlying graphs (weight recovery), as well as reconstruct the full tensor (predictive accuracy). For a first test, we generated a tensor from graphs with varied topologies (including Erdős– Rényi, bipartite, star, and configuration random graph models) to determine whether GTD was flexible enough to recover different types of structures. We also tested GTD on tensors generated from one graph structure at a time (SI S2.7). On this first set of synthetic tests, GTD recovered the underlying graph structure and fit the data well (Fig. 5), demonstrating that GTD is flexible enough to recover structure beyond what its prior assumes.

**Fig. 5.**
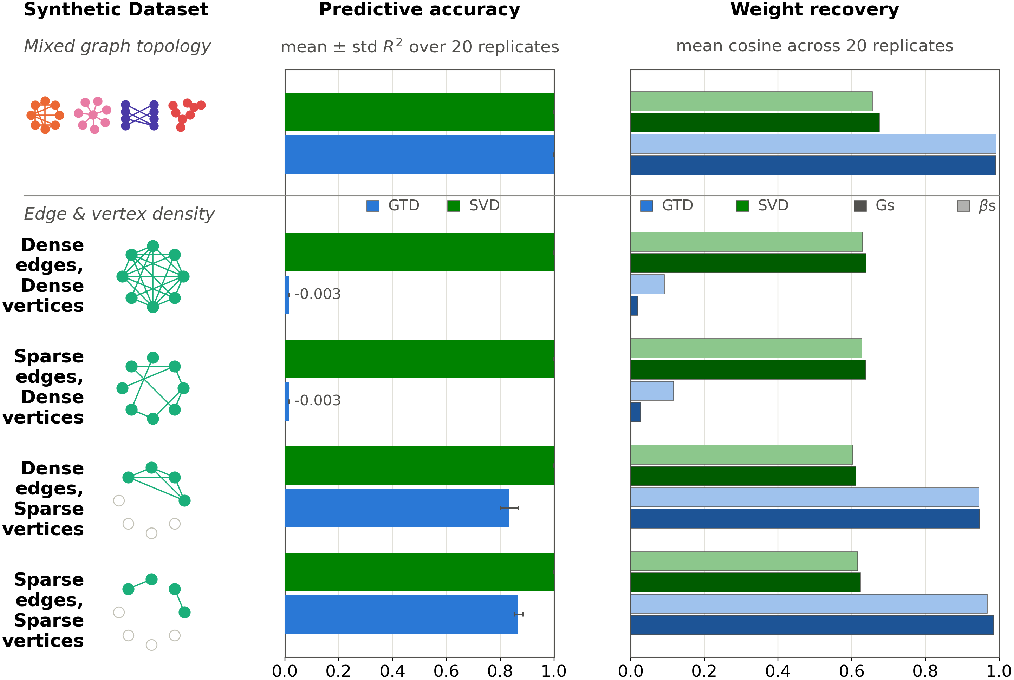
GTD applied to synthetic data. Rows are grouped into two test suites: fixed sparsity with varied graph structure (top), and fixed graph structure (Erdő s–Rényi) with varied sparsity and overlap (bottom). For the first test, we generated *r* = 8 graphs with a support of 10 vertices sampled from *L* = 40 total vertices. For the second, we used *r* = 10 graphs with varied support sizes sampled from *L* = 300 vertices. First column: illustrations of each graph configuration. Second column: mean ± std *R*^2^ for GTD and SVD over 20 independent replicates. This predictive accuracy measures tensor-level reconstruction. Third column: mean ± std cosine similarity to the ground-truth graphs and weights for GTD and SVD over 20 independent replicates.

Because GTD assumes that the basis graphs are sparse, our second set of tests varied the vertex and edge densities under a fixed Erdős–Rényi (ER) random graph model. We generated graphs with substantially overlapping support, both as a more difficult test and a more realistic biological scenario. On this set of tests, GTD performed well with both sparse and dense edge sets. However, GTD was not able to recover structure when we increased the density of vertices (Fig. 5).

For both types of tests, per-environment *β* weights were randomly generated, and we added 10% Gaussian noise relative to the standard deviation of the tensor *T* before running GTD (see SI S2.7 for more details).

We compared our method to the Singular-Value Decomposition (SVD), which flattens *T* and thus cannot exploit tensor structure; this yields components that are typically not sparse, and difficult to interpret biologically. In contrast, GTD penalizes both the number of edges and vertices in its learned components (Eq. 3); it favors edges that share vertices, which is structure that SVD cannot see when it flattens *T* . On synthetic data, SVD found the lowest error solution, but only partially recovered the true generating structure. Hence, low error alone does not guarantee the discovery of sparse, interpretable structure.

We also compared GTD to MCPCA (31). While MCPCA found structure in the *β* weights, it did not recover generating graph structures that were not well-approximated by rank-one matrices (see SI S2.8).

## 3. Application to yeast data

### 3.1. Description of the data

We next applied both SLED and GTD to experimental data from an F1 yeast cross. The data consist of the growth rates of *N* ≈ 100, 000 yeast segregants each measured in *E* = 101 environments (32). The environments have a “hub-and-spoke” structure: each condition consists of a base medium plus a perturbation (stressor). Each perturbation is applied across a range of strengths in each base environment. See Table 1 for a list of base media and perturbations.

**Table 1.** Summary of the 101 environments assayed in the yeast dataset, adapted from (32). The abbreviated base media correspond to yeast extract peptone dextrose, yeast nitrogen base, guanidinium, lithium acetate, and geldanamycin.

|  |  |  | Hub environment |  |  |  |  |
|---|---|---|---|---|---|---|---|
| Perturbation | Cellular processes affected | Gradient range | YPD | YNB | Gu | Li | Gel |
| Temperature | Protein folding, reaction rates | 23 – 37°C | ✓ | ✓ | ✓ | ✓ | ✓ |
| Cycloheximide | Protein synthesis | 50 – 200 nM | ✓ | ✓ | ✓ | ✓ |  |
| Fermentation/Respiration | Energy metabolism | 1.5/1 – 0/4 %glu/gly | ✓ | ✓ | ✓ | ✓ |  |
| Caffeine | DNA repair, TOR pathway | 1 – 8 mM | ✓ | ✓ | ✓ | ✓ |  |
| Glucose | Metabolism, osmotolerance | 6 – 20% | ✓ | ✓ | ✓ | ✓ |  |
| pH | Metabolism, osmotolerance | 2.5 – 8.5 | ✓ |  |  |  |  |
| Sodium chloride | Osmotolerance | 0.1 – 1.2 M | ✓ |  |  |  |  |
| Potassium chloride | Osmotolerance | 0.2 – 1.5 M | ✓ |  |  |  |  |
| 4-nitroquinoline 1-oxide (4NQO) | DNA damage and repair | 10 – 50 ng/ml |  | ✓ |  |  |  |

The segregants are the product of an F1 cross between the BY and RM yeast strains, which differ by ∼ 42,000 single nucleotide polymorphisms (SNPs) (33). Due to the F1 cross, the BY and RM alleles occur with similar frequencies across the segregant panel at most of the SNPs, making this an ideal dataset to search for epistatic structure (34). Likewise, the large number of structured environments are designed to reveal pleiotropic structure. Due to a finite recombination rate, there are an average of just 50 recombination breakpoints along the genome, corresponding to, on average, ∼ 820 SNPs between breakpoints (33). Thus, it is not necessary (nor computationally feasible) to input all the loci into our models. For SLED, we first compressed the genome to a set of 1,429 loci that captured most of the genetic variability of the segregant panel (see SI S1.4.1).

The tensor of epistatic interactions used as the input for GTD was inferred in a previous analysis by a joint forward search procedure (see SI S2.2) (32).

### 3.2. SLED, SSD, and GTD converge on yeast data

We first applied SLED and GTD to the yeast data, then compared the results to those obtained via SSD, which uses only additive information to discover modular structure (23). Comparing these three methods allowed us to analyze how the inferred modular structure depends on the type of genetic information used as input.

In Figs. 6 and 7, respectively, we show SLED and GTD trained on the yeast data. Without any prior knowledge of the environmental structure of the data, both models learned a clearly interpretable, and similar, set of modules, which we order to be consistent across methods. Module 1 is broadly activated, suggesting that it describes regulation and general growth. Modules 2, 3, and 4 are predominantly activated in distinct bases: module 2 in guanidinium, module 3 in lithium, and module 4 in YNB. This is consistent with the lack of a standalone YPD module—unlike the other bases, YPD is not a stressor. For both models, the remaining modules capture more granular, perturbation-specific responses. For SLED, these are module 5 for cycloheximide and modules 6 and 7 for caffeine on different base media. For GTD, these are module 5 for caffeine, module 6 for cycloheximide, module 7 for YNB 4NQO, and module 8 for a subset of the lithium environments.

**Fig. 6.**
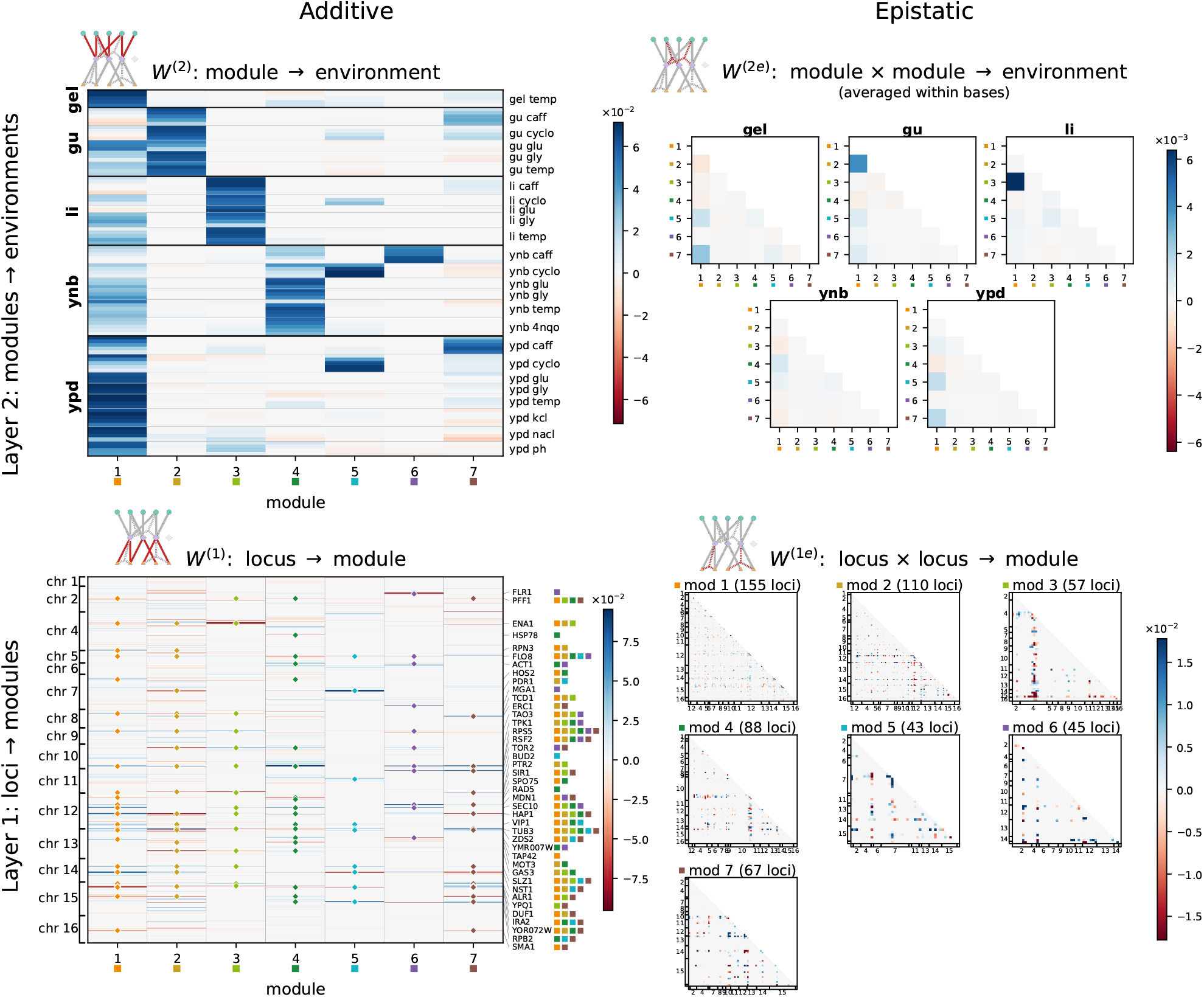
SLED applied to a dataset of the growth rates of yeast segregants in 101 environments with a hub-and-spoke structure. Top left: additive module-to-environment map. Base environments are separated by thick black lines; perturbations within a base are separated by thin gray lines. Top right: epistatic module-to-environment map. The epistatic interactions between modules, averaged within each of the five base environments. Bottom left: additive locus-to-module map. Strong effect loci are annotated, with colored squares indicating the modules they participate in. If a locus is annotated, other nonzero effects of that locus in other modules are also annotated. Bottom right: epistatic locus-to-module map. The epistatic interactions between loci in each module, with the heatmaps filtered to show only the participating loci.

**Fig. 7.**
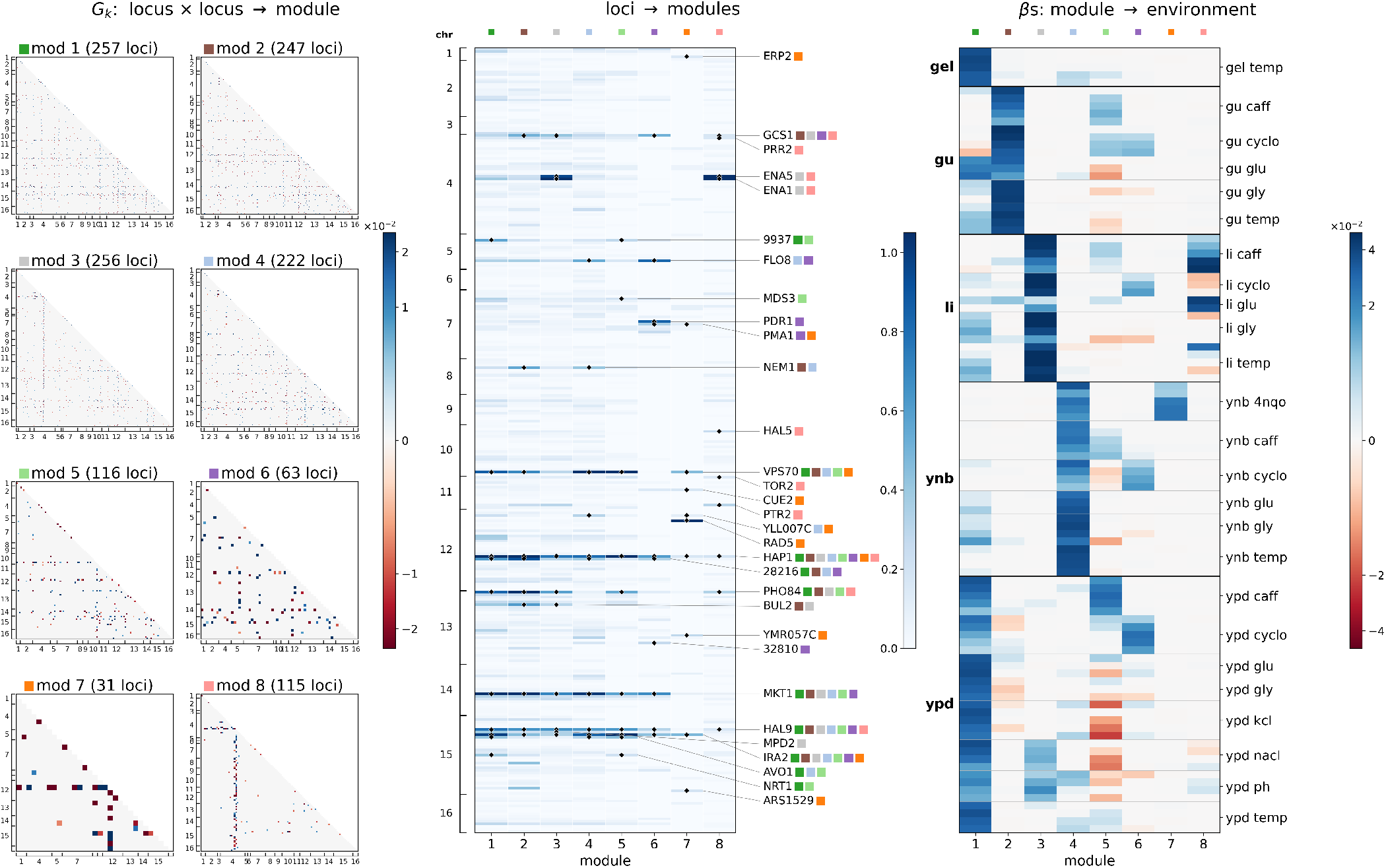
GTD applied to the inferred epistatic weights from the yeast dataset. Left: epistatic locus-to-module map. An interaction is included in a module if it is stronger than a null baseline (see SI S2.6). Loci are labeled by chromosome. Middle: strength of loci within a module (weighted sum of interaction magnitudes). The top 10 strongest loci in each module are annotated, along with a colored square that indicates the module(s) they are active in. Right: module-to-environment map; base environments are separated by thick black lines, and perturbations within them by thin gray lines.

The clear separation of the modules strongly suggests that both SLED and GTD capture the relevant genetic features that determine fitness in each condition. For example, the known regulatory genes FLO8 (35), IRA2 (36), and HAP1 (37) participate in module 1 of both models. Additionally, modules corresponding to a stressor each include genes that have known roles in tolerating that stressor. For SLED, PDR1 (38) participates in modules 2 and 5 (guanidinium and cycloheximide), ENA1 (33, 39) in module 3 (lithium), and FLR1 and ERC1, which encode drug/toxin efflux transporters (40– 42), separately in modules 6 and 7 (caffeine). For GTD, ENA1 and HAL5 (39) participate in modules 3 and 8 (lithium), PDR1 in module 6 (cycloheximide), RAD5 (43) in module 7 (YNB 4NQO), and MDS3 (44, 45) in module 5 (caffeine). Across stressors, the recovered genes span distinct tolerance mechanisms, including efflux (ENA1 and PDR1 (39, 46)), ion homeostasis (HAL5 (47)), the drug’s cellular target pathway (MDS3 (48)), and DNA repair (RAD5). Interestingly, the separation of modules 6 and 7 by SLED suggests that caffeine acts differently on the YNB base than on the guanidinium, lithium, or YPD bases. At the genetic level, the two modules have only partially overlapping gene sets. The fact that SSD (see below) and SLED make a distinction between caffeine on YNB versus other bases, but GTD does not, suggests that the differences appear in the additive effects, rather than emerging from unique patterns of epistatic interactions that differ between the two bases. Finally, we note that both methods also identify several putative genes of unknown function with effects across environments, e.g. YOR072W for SLED and YLL007C for GTD.

We next examine the patterns of epistasis learned by each model. For both models, the structure of intra-modular epistasis varies significantly across modules. For example, epistasis within modules 1 and 2 is broad and diffuse, with interactions distributed across the genome. In contrast, specialized modules are primarily organized around individual hub loci. Epistasis within module 3, for example, is dominated by interactions with ENA1, which also has the strongest additive effect in that module. Thus, our models capture both broad, genome-wide patterns of epistasis as well as strong, hub-specific effects, consistent with the known biology of yeast (49).

SLED allows us to examine two features of epistasis that GTD does not. First, SLED infers patterns of inter-modular epistasis. Intuitively, we find this to be dominated by interactions between the general module (module 1) and modules specific to a relevant base environment. Particularly strong interactions occur between the general module and the guanidinium (module 2) and lithium (module 3) modules. Across the base environments, lithium exhibits the strongest signatures of inter-modular epistasis, while YNB is the most additive.

Second, by disabling one or both kinds of epistasis in SLED, we can estimate the relative importance of intra- (between genes within a module) versus inter-modular (between modules) epistasis. Disabling intra-modular epistasis, inter-modular epistasis, or both yielded, respectively, 6%, 0.5%, and 7% higher MSE on average, compared to full SLED models with comparable regularization. This agrees with previous work: epistatic interactions make a small but significant contribution to predictive accuracy, often involving at least one locus with a strong additive effect (7). Finally, intra-modular epistasis appears to be more important than inter-modular epistasis. More concretely, in this dataset, we find that groups of loci that have coordinated additive effects are more likely to exhibit epistasis within, rather than across, such groups.

Returning to the comparison between SLED and GTD, we examine the pleiotropic structure learned by each model. In both models, many of the strongly activated loci participate in multiple modules and thus are, to varying degrees, pleiotropic. Most of these pleiotropic genes participate in module 1, consistent with the idea that perturbation-specific responses modify a general regulatory program. We also see pleiotropic behavior in regulatory genes (e.g. FLO8, HAP1, and IRA2) and genes that function in stress response (e.g. ENA1), as identified by both methods. Intriguingly, for SLED, some modules contribute positively in some environments but negatively in others, which indicates groups of genes for which the beneficial allele is not constant across environments. This is also true for GTD with respect to whether a combination of genes is better or worse than the sum of its parts in a given environment. Overall, the qualitative pattern of pervasive pleiotropy, concentrated in a general growth program and in stress-response hubs, is in line with previous characterizations of the genetic architecture of yeast (9, 33).

Next, we compare the genetic architectures inferred by SLED, SSD, and GTD. Despite the differences between the models with respect to the type of input information (additive, epistatic, or both), model architecture, regularization, training procedure, and even set of input loci, the genetic and modular structures discovered by the three methods are highly similar. Across the models, six modules converge (out of seven total for SLED, ten for SSD, and eight for GTD) (Fig. 8b). The differences are as follows: SSD and GTD learn a module for 4NQO on YNB; SLED and SSD learn a module for caffeine on YNB; GTD learns a lithium sub-module; and SSD learns two more subtle modules that SLED and GTD do not. We view these differences largely as refinements of the convergent structure, e.g. additional environments folded into an existing module, or a module splitting into smaller sub-modules. This also explains why the total number of modules varies across the three methods.

**Fig. 8.**
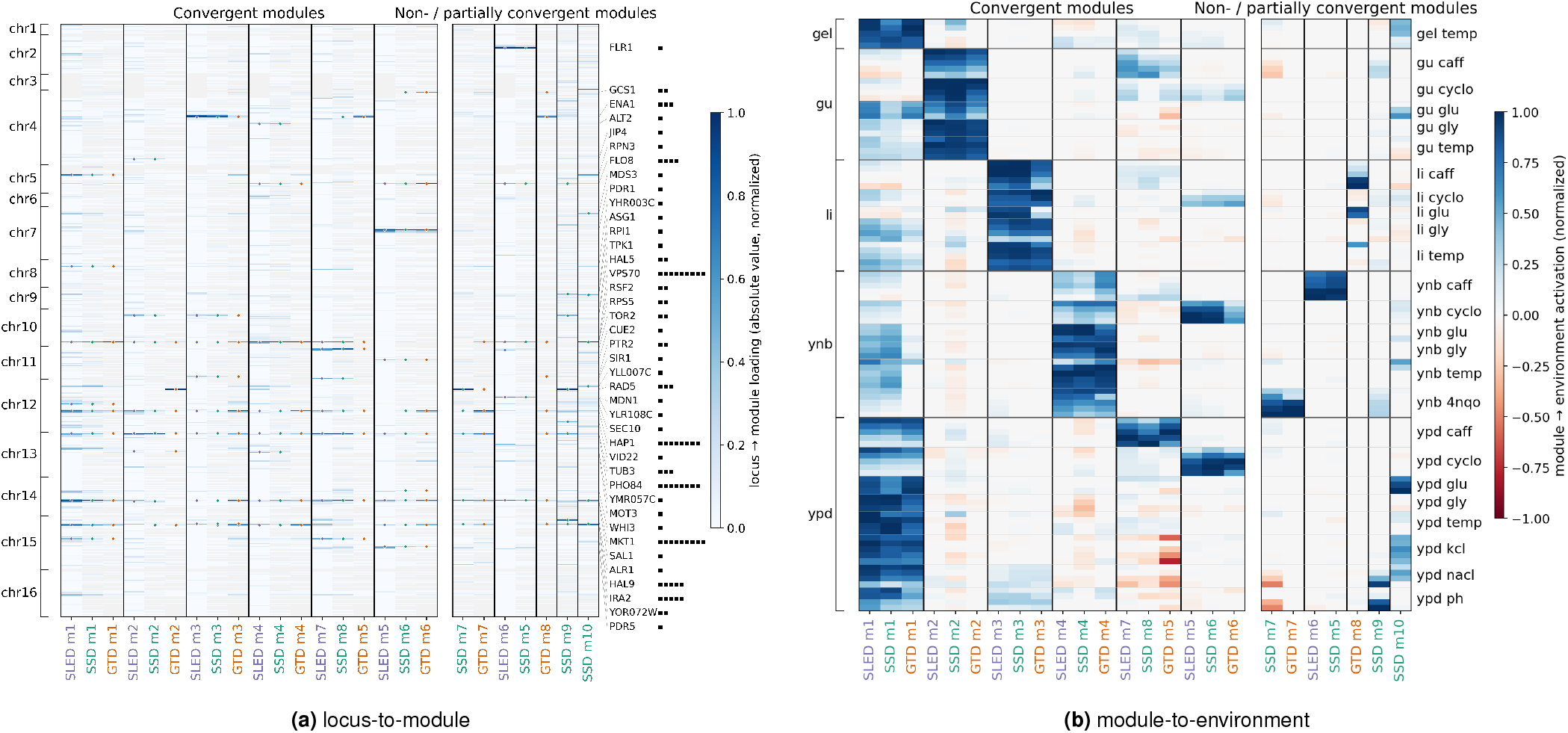
Comparison of modular structure inferred by SLED, SSD, and GTD. **(a)** For each model, convergent modules are aligned based on their environmental activation profiles (right). The additive locus-to-module maps for each method are shown, grouped by corresponding modules. The weights are reported in absolute value and normalized relative to the largest effect size within each module. The models have different input gene sets (SLED uses 1,429 loci; SSD 382; and GTD 308), so genes are binned in 20 kb intervals for alignment. Gray indicates genetic regions not used as inputs to a particular model. Select strong effect genes are annotated, as in Figs. 6 and 7. Black squares indicate how many modules a locus participates in. **(b)** For each model, the additive module-to-environment maps are shown, grouped by convergent modules. The activations are normalized to the maximum within each module. In both panels, within a convergent module cluster, SLED is leftmost, SSD is in the center, and GTD is rightmost.

Many of the genes involved in each of the six convergent modules are also highly consistent across the three models (Fig. 8a). For SSD and GTD, this is partially to be expected due to their shared pre-processing (see SI S2.2). However, this pre-processing does not guarantee the highly correlated modular partitioning that we observe, and, furthermore, SLED converges with SSD and GTD despite using an expanded locus set. On a more granular level, the methods are particularly similar in their treatment of strong effect and highly pleiotropic loci, such as ENA1 in the lithium base module. Nonetheless, as expected, the module-to-environment map is more consistent than the locus-to-module map, since resolving specific causal loci is harder than identifying the environmental effect they jointly produce.

To summarize, the convergent modularity discovered by these three different methods suggests that pleiotropic additive and epistatic effects point to the same underlying genetic architecture governing the variation in growth rates across environments.

### 3.3. Generalization to novel environments

To validate the modular structure discovered by each model, we tested the abilities of both SLED and GTD to generalize to novel environments. We hypothesized that, if the models learned modules that approximate the true genetic architecture, these pre-inferred modules could be used to predict phenotypes in novel environments.

The structure of the yeast data naturally lends itself to a set of increasingly challenging generalization tests. We can hold out from the training data a random subset of environments, a perturbation on a single base medium (or “block”), a perturbation across all base media, or an entire base medium. These tests add layered demands for the models because they ask them to generalize to increasingly foreign environments. We expected the models to generalize well to environments that are “similar” to the training environments, and to fail on environments that are “distant” from the training environments, though it is difficult to make this notion of environmental similarity precise.

For both SLED and GTD, we first trained models with groups of environments held out, then retrained a subset of parameters to fit these environments while keeping the rest fixed. Specifically, for SLED, we froze the first-layer weights, encoding the organism’s genetic architecture, and retrained only the second-layer weights, which determine which features of that architecture are important in any given environment. For GTD, we froze the learned graphs and retrained the perenvironment *β* weights on all of the environments. We view the graphs as generalizable epistatic patterns, with their activations adjusted in specific environments. In both cases, we quantified the ability of each model to generalize by comparing the test MSE of each held-out model to a model trained on all environments (Fig. 9).

**Fig. 9.**
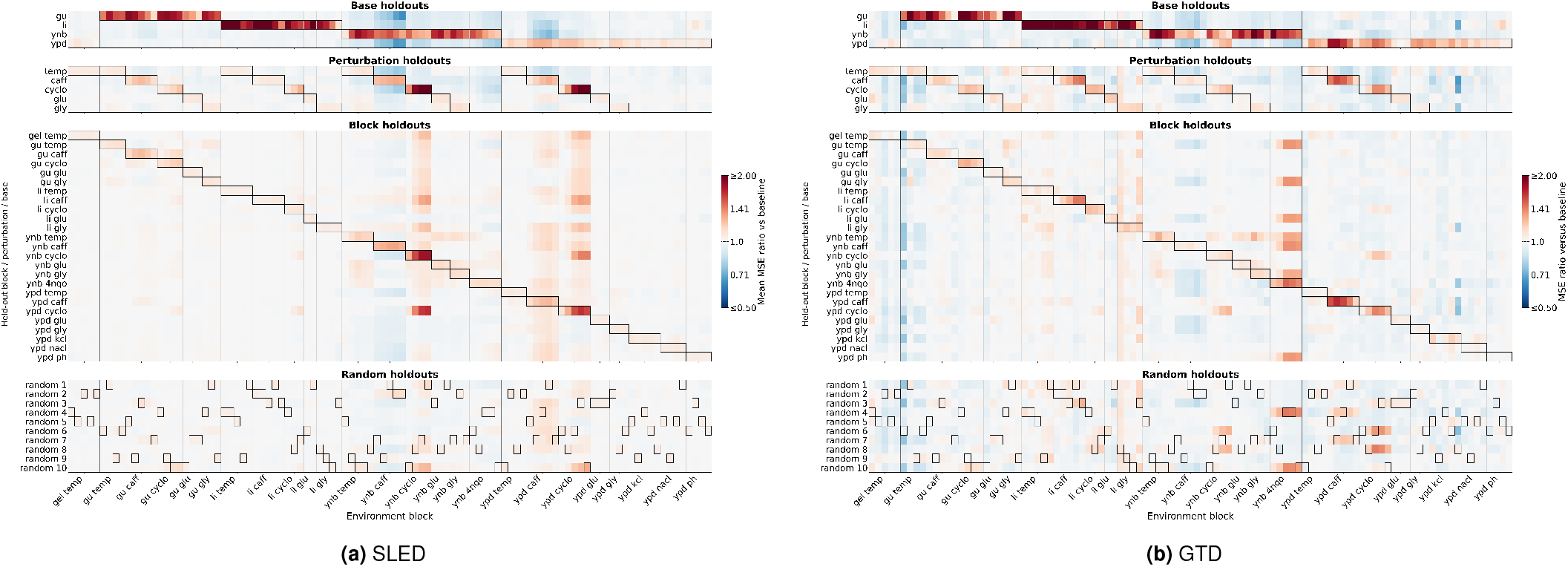
Generalization of SLED and GTD to novel environments. For both panels, rows are grouped into the four types of held-out tests: a base medium, a perturbation across base media, a perturbation within a single base medium (“block”), and random hold-outs. Within groups, each row represents one such test configuration. Each random test is a disjoint set of 10 environments (10% of the total). The environments held out by a given configuration are outlined in black. The color bar indicates the ratio of the MSE of each row configuration to a baseline model trained on all environments. **(a)** For SLED, the MSE is computed between the predicted and observed phenotypes. The reported values are averaged pairwise over replicate fits: 50 baseline and 10 of each hold-out. **(b)** For GTD, the MSE is the reconstruction error of the inferred weight tensor. GTD is deterministic, so there is no averaging.

The generalization capabilities of the two models are quite similar. As expected, the models generalize well on random and single block hold-outs, presumably because the held-out environments are similar to environments in the training set. Performance degrades when entire base media are held out (in part due to the large decrease in training data) or, to a lesser extent, when an entire perturbation is held out. This is intuitive: for example, fitness in the lithium base is strongly determined by the ENA1 allele, the effect of which the model does not learn when the lithium environments are omitted from the training data. This example underscores the fact that what *any* model learns about the GP map is highly dependent on the set of experimental conditions analyzed. Overall, these results demonstrate the strong generalization capabilities of our models, suggesting that the modular organization that our models learn accurately captures the underlying biological structure.

## 4. Discussion

Here, we introduced two complementary methods for identifying interpretable modular structure in multi-trait, whole-genome GP maps. The first, SLED, infers modules directly from GP data and models both additive and epistatic effects. The second method, GTD, identifies modules from a pre-inferred tensor of epistatic effects. Both methods assume that the GP map has a sparse, low-dimensional structure, with core genetic modules present across subsets of environments. On synthetic data, we showed that both methods reliably recovered known underlying structure and did not impose such structure in its absence. We then applied both methods to data consisting of the relative growth rates of F1 segregants from a yeast cross measured across structured environments. Both methods demonstrated a strong capacity to generalize by successfully predicting relative growth rates in novel environments using pre-learned modules. Finally, we compared the outputs of SLED, SSD (a previous method that does not model epistasis), and GTD to explore how the inferred modular architectures depended on the type of genetic information available to a model. We found that, on this dataset, the three methods yielded very similar modular structures.

The convergent findings of SLED, SSD, and GTD suggest that modular organization is a useful coarse view of the GP map analyzed here. Importantly, this convergence is not guaranteed *a priori*. SLED assumes a shared modular structure between additive and epistatic effects, which we explore in more detail below. GTD’s modules could, in theory, diverge entirely from the modules inferred from first-order pleiotropic effects. While the pre-processed *G* × *G* × *E* epistatic effects tensor we input into GTD only includes interactions among loci with inferred additive effects, it imposes no constraint on how these loci should be grouped into modules. As such, the agreement between the structure inferred by SLED and GTD suggests that patterns of additive and epistatic effects are similar, but not simply as an artifact of shared modeling assumptions.

Of course, this convergence depends on the depth of the dataset. We do not have any power to resolve modules that, hypothetically, do not vary across the genotypes in the dataset or the environments in which we measure phenotypes. Moreover, we also may not be able to detect small effect loci, and their patterns of interaction may not fit cleanly into those found here. For this reason, the most complete picture of the GP map will emerge from data that probe as many genotypes and distinct environments as possible. However, it is worth noting that SLED converges on this dataset even when restricted to approximately one-third of the data (*N* ∼ 30, 000 genotypes, see SI S1.4.3).

SLED and GTD complement a growing family of models of GP maps. Methods such as Epistatic Factor Analysis, D-LIM, and MAVE-NN, as well as Gaussian process methods including LANTERN and product kernels, capture epistasis but are designed for a single phenotype and thus do not account for pleiotropy (50–54). Conversely, current methods that extend to multiple phenotypes are constrained by their inputs or prior assumptions: MoCHI relies on specified biophysical models and is tailored for deep mutational scanning data, and GenNet requires prior gene annotations or pathway information (55, 56). Finally, recent attempts to apply machine learning methods to GP maps have yielded improvements in predictive accuracy, but at the expense of interpretability (24). SLED and GTD are therefore distinguished by inferring interpretable modular structure jointly from patterns of epistasis and pleiotropy.

SLED imposes several structural constraints on the form of inferred epistasis. First, due to its linked-epistasis prior, SLED will fail to capture epistatic interactions between loci without marginal additive effects. Second, SLED allows epistasis only within and between modules, which prohibits interactions between loci whose additive effects are decoupled across environments. Third, when two modules have an epistatic interaction, then all the genes within the two modules implicitly have an epistatic interaction. Finally, SLED cannot capture interactions beyond fourth-order in loci, nor can it explicitly model global epistasis.

Additionally, because the number of possible epistatic interactions scales quadratically with the number of input loci, SLED is computationally feasible only for datasets containing up to approximately 10^4^ loci (see Fig. S1 for a quantitative runtime analysis). However, as in this paper, it is often possible to pre-process long genotypes into a smaller set of loci that capture the majority of genotypic variance.

GTD, in turn, is constrained by the structure of its input tensor. In this paper, the input *G* × *G* × *E* tensor of epistatic effects is substantially sparse (32) and GTD partitions this signal into interpretable subgroups. However, since GTD assumes that loci group into a small number of sparse, shared graphs, it is not designed to recover truly dense structure where all loci participate in every module (see SI S2.7).

These design choices determine when each method is preferable. SLED acts directly on GP data and jointly models additive and epistatic effects, but assumes that these effects share the same modular structure. Future work could apply SLED to datasets for which this assumption is plausible, including human phenotype, RNA-sequencing, cell-line, protein, and microbial ecology datasets. GTD, in contrast, linearly decomposes epistatic effects in order to recover modular structure from patterns of epistasis across environments. This method is therefore ideal for datasets that primarily characterize epistasis, such as double-deletion collections in model organisms and proteins, or high-powered assays for which pairwise epistasis can be reliably inferred, as demonstrated here.

Across disciplines, there is a trade-off between accuracy and interpretability in model development (57, 58). While models of GP maps are often evaluated based on their predictive accuracy, an equally important goal is to generate hypotheses about the structure of genetic architecture, both providing insight into principles of biological organization and guiding experimental design. Exploring datasets with a variety of interpretable models is thus imperative when comparing and testing hypotheses about the nature of biological organization.

## Acknowledgments

S.N.P. is supported by the Burroughs-Wellcome Fund’s Career Awards at the Scientific Interface (CASI). M.M.D is supported by NIH grants R01-GM104239 and R35-GM161612. C.M.H is supported by the Jane Coffin Childs Memorial Fellowship. A.S. is supported by NSF DMS 2608217 and an Alfred P. Sloan research fellowship. The authors acknowledge the Tufts University High Performance Compute Cluster (https://it.tufts.edu/high-performance-computing) and the FASRC Cannon cluster supported by the FAS Division of Science Research Computing Group at Harvard University (https://www.rc.fas.harvard.edu).

