## Supplemental Information for "Inferring modularity in genotype-phenotype maps using patterns of epistasis and pleiotropy"

##### This PDF file includes:

Supporting text
Figs. S1 to S11
Tables S1 to S2
SI References

### Supporting Information Text

#### S1. Supplementary text for SLED

##### S1.1. Model training and selection.

***S1.1.1. Model definition.*** SLED is a model for predicting phenotypes from genotypes via a sparse, low-dimensional map that incorporates epistasis, both between genes in a module and between modules. Explicitly,

$$m_n = \sum_{j=1}^{L} W_{nj}^{(1)} g_j + \frac{1}{2} \sum_{j,k=1}^{L} W_{njk}^{(1e)} g_j g_k + b_n \quad \text{[S1]}$$

$$\hat{p}_e = \sum_{n=1}^{M} W_{en}^{(2)} m_n + \frac{1}{2} \sum_{n,m=1}^{M} W_{enm}^{(2e)} m_n m_m \quad \text{[S2]}$$

where the weights $W$ are encouraged to be sparse through regularization. As discussed below, we z-score the phenotypes before fitting the model, and thus omit the bias in the second layer.

***S1.1.2. Pre-processing.*** Minimal pre-processing is required before model training. We z-score the phenotypes such that the mean phenotype in each environment is zero. The variance is an arbitrary choice; we found $\sigma^2 = 0.01$ yielded good numerical stability. Z-scoring the phenotypes ensures that each environment is given equal weight in the loss function and removes the need for a bias—roughly, the mean phenotype in each environment—in the second layer. It is straightforward to convert back to the original units if desired.

Additionally, we mean center the genotypes, i.e., we send $g_j \to g_j - \langle g_j \rangle$, where the brackets denote a population average. We do this because, when the probability of finding a particular allele across the population is not 1/2, the asymmetry of the allele distribution can induce an effective additive weight from an epistatic interaction. To see this, consider the expression for module $n$

$$m_n = \sum_{j} W_{nj}^{(1)} g_j + \frac{1}{2} \sum_{j,k} W_{njk}^{(1e)} g_j g_k + b_n \quad \text{[S3]}$$

Writing $g_j = \langle g_j \rangle + \delta g_j$, where the angle brackets denote a population average, we have

$$m_n = \sum_{j} \left[ W_{nj}^{(1)} + \sum_{k} W_{njk}^{(1e)} \langle g_k \rangle \right] g_j + \frac{1}{2} \sum_{j,k} W_{njk}^{(1e)} \delta g_j \delta g_k - \frac{1}{2} \sum_{j,k} W_{njk}^{(1e)} \langle g_j \rangle \langle g_k \rangle + b_n \quad \text{[S4]}$$

where the coefficient in front of the epistatic contribution to the effective additive weight is 1 instead of 1/2 due to the symmetry of the epistasis tensor in locus indices. For this reason, we mean-center the genotypes before training to prevent the model from inducing an additive effect via an epistatic weight, as in the first term of Eq. (S4). When $\langle g_j \rangle = 0$, it is easy to see that the expression simplifies as expected.

***S1.1.3. Model training.*** We fit the weights $W^{(1)}, W^{(1e)}, W^{(2)},$ and $\mathrm{W}^{(2e)}$ using standard stochastic gradient descent with an Adam optimizer on the following loss function:

$$\mathcal{L} = \frac{1}{NE} \sum_{i=1}^{N} \sum_{e=1}^{E} \left( p_e^{(i)} - \hat{p}_e^{(i)} \right)^2 \quad \text{[S5]}$$

$$+ \lambda_1 \sum_{n=1}^{M} \sum_{j=1}^{L} \log \left( 1 + \left| W_{nj}^{(1)} \right| \right) + \lambda_2 \sum_{e=1}^{E} \sum_{n=1}^{M} \log \left( 1 + \left| W_{en}^{(2)} \right| \right) \quad \text{[S6]}$$

$$+ \lambda_{1e} \sum_{j,k=1}^{L} \sum_{n=1}^{M} \frac{\log \left( 1 + \left| W_{njk}^{(1e)} \right| \right)}{\left| W_{nj}^{(1)} W_{nk}^{(1)} \right| + \epsilon} + \lambda_{2e} \sum_{n,m=1}^{M} \sum_{e=1}^{E} \frac{\log \left( 1 + \left| W_{enm}^{(2e)} \right| \right)}{\left| W_{en}^{(2)} W_{em}^{(2)} \right| + \epsilon} \quad \text{[S7]}$$

where the superscript in the first line denotes individual $i$. The first term in the loss is the mean-squared error (MSE) between the predicted and observed phenotypes across environments. The second and third terms encourage the additive weights to be sparse. For small weights, these regularizers are similar to an L1 norm: $\log(1 + |\epsilon|) = |\epsilon| + \mathcal{O}(\epsilon^2)$. For large values of the weights, the regularization is nearly constant with respect to the magnitude of the weight. Roughly speaking, we penalize the existence but not the magnitude of the weights. The numerators of the last two terms are identical norms on the epistatic weights, but these are inversely weighted by the additive contributions of the loci (or modules) that are to be linked epistatically. This means that epistatic weights are penalized by a factor of at least $1/\epsilon$ (where $\epsilon = 0.01$) unless both loci already contribute to the same module (and similarly from module to phenotype), as desired. The four hyperparameters $\{\lambda_1, \lambda_{1e}, \lambda_2, \lambda_{2e}\}$ control the relative contribution of each term to the total loss, and thus the degree of sparsity of each type of interaction. Also note that the diagonals of the epistatic tensors are zero, i.e. $W_{njj}^{(1e)} = W_{enn}^{(2e)} = 0$ for all $j, n$, prohibiting self-interactions. Additionally, the interaction tensors are symmetric in the appropriate indices, i.e. $W_{njk}^{(1e)} = W_{nkj}^{(1e)}$, $W_{enm}^{(2e)} = W_{emn}^{(2e)}$.

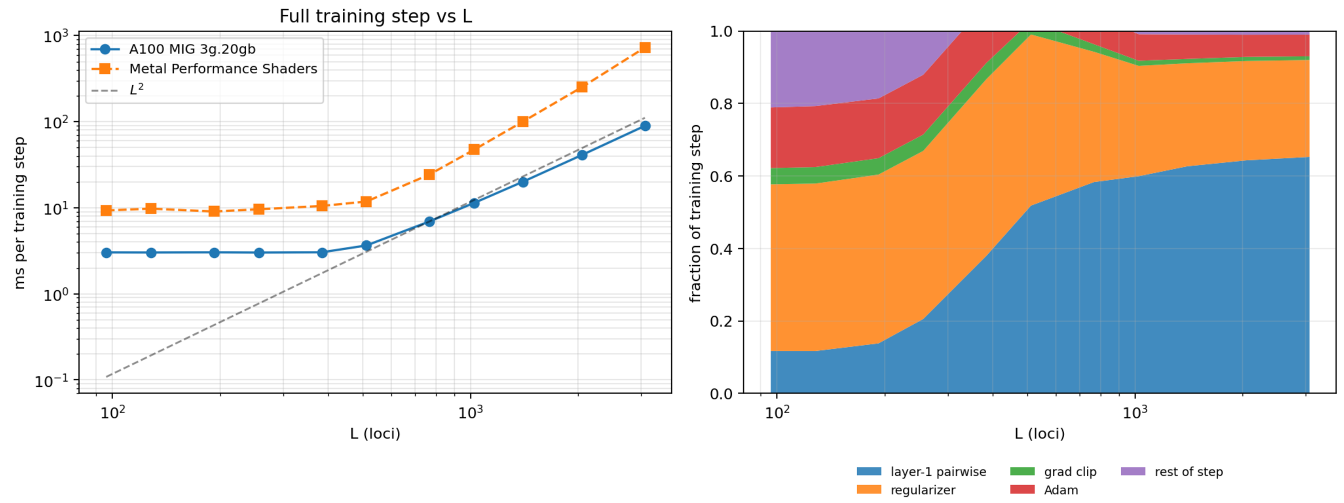


**Fig. S1.** Runtime scaling with the number of loci L, at fixed $M = 10$, $E = 101$, and minibatch size $B = 1024$. (Left) Wall-clock time per training step on an NVIDIA A100 (MIG 3g.20gb) and an Apple M5 Pro GPU, with a dashed quadratic reference. (Right) Fraction of the training step spent on each component on the A100.

***S1.1.4. Runtime scaling analysis.*** In Fig. S1, we analyze the scaling of SLED runtime with input genotype length ($L$). We measured time per training step as a function of the number of loci, holding the module count ($M = 10$), environment count ($E = 101$), and minibatch size ($B = 1024$) fixed. The number of epistatic parameters grows exactly as $L^2$, but runtime does not: on an NVIDIA A100 GPU (MIG 3g.20gb partition), time per step scaled as $L^{1.63}$ over $L = 384$–$3072$. In practical terms, one epoch over 45,000 training genotypes (43 minibatches of size 1024) took 0.13 s at $L = 384$ and 0.86 s at $L = 1400$, so a 1000-epoch run at $L = 1400$ completes in roughly 14 minutes. For comparison, the same sweep on an Apple M5 Pro GPU scaled as $L^{2.04}$.

***S1.1.5. Model selection.*** We first input an upper bound on the number of modules, in this case choosing 10. However, the regularization (described above) forces some of the modules to zero, so the effective number of modules is not independent of the regularization strength. The model discussed in the main text had 7 nonzero modules.

We swept the four sparsity hyperparameters in log space, each ranging from $10^{-8}$ to $10^{-2}$. This yielded a four-dimensional tensor of the MSE on the validation set as a function of the four hyperparameters. The MSE increases monotonically with each of the hyperparameters, rising from some minimum value for a weakly regularized model, to the variance of the phenotypes for a very strongly regularized model that effectively learns the mean phenotype in each environment. In practice, there exists a four-dimensional analogue of an "elbow," where the MSE begins to rise significantly as the model becomes more strongly regularized.

By choosing a contour of constant MSE in this four-dimensional space, one can examine a family of models that trade off sparsity in different regimes of the model but maintain the same predictive accuracy. We emphasize that there is no principled way of choosing between models with equal error; they emphasize different kinds of interactions. The model we present in the main text is one such choice of model, chosen to balance the sparsity in each of the sets of weights roughly evenly.

The models presented in the main text (with $L = 1,429$ and $E = 101$) use the following hyperparameters: $M = 10$, $\lambda_1 = 2.55 \times 10^{-6}$, $\lambda_2 = 6.00 \times 10^{-4}$, $\lambda_{1e} = 1.64 \times 10^{-7}$, and $\lambda_{2e} = 3.98 \times 10^{-5}$.

***S1.1.6. Model weight degeneracy.*** There exists a continuous degeneracy of the weights under which the phenotypic prediction matrix (but not the regularization term in the loss function) is invariant. The degeneracy can be represented by the following transformation:

$$W_{nj}^{(1)} \to \frac{W_{nj}^{(1)}}{c_n}, W_{njk}^{(1e)} \to \frac{W_{njk}^{(1e)}}{c_n}, b_n \to \frac{b_n}{c_n} \quad \text{[S8]}$$

$$W_{en}^{(2)} \to c_n W_{en}^{(2)}, W_{enm}^{(2e)} \to c_n c_m W_{enm}^{(2e)} \quad \text{[S9]}$$

where $c_n$ is a per-module constant. In order to make the weights directly comparable across modules, after training we choose $c_n$ such that, for each module $m_n$, the distribution of module values across the population has unit variance. We also fix the sign of the modules (whether $c_n$ is positive or negative) by requiring that the majority of the activation weight for each module be positive. This rescaling is purely a matter of interpretation and does not affect the predictions of the model. Importantly, breaking this degeneracy means that both the signs and magnitudes of weights can be directly compared across modules.

#### S1.2. SLED as a tensor decomposition algorithm.

***S1.2.1. Effective weights.*** Since SLED is a map from genotype to phenotype, it defines a set of "global" effective weights that map genotype directly to phenotype. These can be obtained by marginalizing over the intermediate modules. Writing, for a single genotype to lighten the notation,

$$\hat{p}_e = \sum_j W^{\text{lin,eff}}_{ej} \delta g_j + \frac{1}{2} \sum_{j,k} W^{\text{epi,eff}}_{ejk} \delta g_j \delta g_k \quad \text{[S10]}$$

by marginalizing over the latent modules, and where we have made explicit that the weights act on mean-centered genotypes. The effective weights can be written in terms of the four sets of learned weights and the empirical one-, two-, and three-point functions of the genotypes. Writing the full expression for a predicted phenotype in terms of its genotype and grouping by order in the genotype, we obtain

$$\begin{aligned}
\hat{p}_e = &\sum_n \left[ W^{(2)}_{en} + \tfrac{1}{2} \sum_m W^{(2e)}_{enm} b_m \right] b_n \\
&+ \sum_j \left[ \sum_n W^{(2)}_{en} W^{(1)}_{nj} + \tfrac{1}{2} \sum_{n,m} W^{(2e)}_{enm} W^{(1)}_{mj} b_n + \tfrac{1}{2} \sum_{n,m} W^{(2e)}_{enm} W^{(1)}_{nj} b_m \right] g_j \\
&+ \sum_{j,k} \left[ \tfrac{1}{2} \sum_n W^{(2)}_{en} W^{(1e)}_{njk} + \tfrac{1}{2} \sum_{n,m} W^{(2e)}_{enm} W^{(1)}_{nj} W^{(1)}_{mk} + \tfrac{1}{4} \sum_{n,m} W^{(2e)}_{enm} W^{(1e)}_{mjk} b_n + \tfrac{1}{4} \sum_{n,m} W^{(2e)}_{enm} W^{(1e)}_{njk} b_m \right] g_j g_k \\
&+ \sum_{j,k,l} \left[ \tfrac{1}{4} \sum_{n,m} W^{(2e)}_{enm} W^{(1)}_{nj} W^{(1e)}_{mkl} + \tfrac{1}{4} \sum_{n,m} W^{(2e)}_{enm} W^{(1e)}_{njk} W^{(1)}_{ml} \right] g_j g_k g_l \\
&+ \frac{1}{8} \sum_{n,m,j,k,l,r} W^{(2e)}_{enm} W^{(1e)}_{njk} W^{(1e)}_{mlr} g_j g_k g_l g_r
\end{aligned} \quad \text{[S11]}$$

where we have grouped the terms by their order in loci. It is easiest to read each term from right to left, imagining a diagrammatic construction in linear or pairwise contributions to each phenotype. We now marginalize over the higher order interactions to obtain the global effective weights, finding

$$\begin{aligned}
W^{\text{lin,eff}}_{ej} = &\sum_n W^{(2)}_{en} W^{(1)}_{nj} + \sum_{n,m} W^{(2e)}_{enm} W^{(1)}_{mj} b_n \\
&+ \sum_k \Bigg[ \sum_n W^{(2)}_{en} W^{(1e)}_{njk} + \sum_{n,m} W^{(2e)}_{enm} W^{(1e)}_{mjk} b_n \\
&\quad + \tfrac{1}{2} \sum_{n,m} W^{(2e)}_{enm} W^{(1)}_{nj} W^{(1)}_{mk} + \tfrac{1}{2} \sum_{n,m} W^{(2e)}_{enm} W^{(1)}_{nk} W^{(1)}_{mj} \Bigg] \langle g_k \rangle \\
&+ \sum_{k,l} \Bigg[ \tfrac{1}{4} \sum_{n,m} W^{(2e)}_{enm} W^{(1)}_{nj} W^{(1e)}_{mkl} + \tfrac{1}{2} \sum_{n,m} W^{(2e)}_{enm} W^{(1)}_{nk} W^{(1e)}_{mjl} \\
&\quad + \tfrac{1}{4} \sum_{n,m} W^{(2e)}_{enm} W^{(1e)}_{nkl} W^{(1)}_{mj} + \tfrac{1}{2} \sum_{n,m} W^{(2e)}_{enm} W^{(1e)}_{njk} W^{(1)}_{ml} \Bigg] \langle g_k g_l \rangle \\
&+ \sum_{k,l,r} \Bigg[ \tfrac{1}{4} \sum_{n,m} W^{(2e)}_{enm} W^{(1e)}_{njk} W^{(1e)}_{mlr} \\
&\quad + \tfrac{1}{4} \sum_{n,m} W^{(2e)}_{enm} W^{(1e)}_{nkl} W^{(1e)}_{mjr} \Bigg] \langle g_k g_l g_r \rangle
\end{aligned} \quad \text{[S12]}$$

Similarly, the effective epistatic tensor is given by

$$\begin{aligned}
W^{\text{epi,eff}}_{ejk} = &\sum_n W^{(2)}_{en} W^{(1e)}_{njk} + \sum_{n,m} W^{(2e)}_{enm} W^{(1)}_{nj} W^{(1)}_{mk} + \sum_{n,m} W^{(2e)}_{enm} W^{(1e)}_{mjk} b_n \\
&+ \sum_l \left[ \sum_{n,m} W^{(2e)}_{enm} W^{(1)}_{nl} W^{(1e)}_{mjk} + \sum_{n,m} W^{(2e)}_{enm} W^{(1)}_{nj} W^{(1e)}_{mkl} + \sum_{n,m} W^{(2e)}_{enm} W^{(1)}_{nk} W^{(1e)}_{mjl} \right] \langle g_l \rangle \\
&+ \sum_{l,r} \left[ \tfrac{1}{2} \sum_{n,m} W^{(2e)}_{enm} W^{(1e)}_{njk} W^{(1e)}_{mlr} + \sum_{n,m} W^{(2e)}_{enm} W^{(1e)}_{nlk} W^{(1e)}_{mjr} \right] \langle g_l g_r \rangle
\end{aligned} \quad \text{[S13]}$$

On a technical note, in all averaged contractions, the summation indices are restricted to loci not in the output set. For example, for a term contributing to $W^{\text{lin,eff}}_{ej}$, any averaged index $\mu$ runs over $\mu \neq j$.

The effective additive weights have contributions from the one-, two-, and three-point functions of the loci, while the epistatic weights, by construction, can only pick up contributions from the one- and two-point functions. Both expressions simplify if we mean-center the genotypes, such that $\langle g_j \rangle = 0$. In practice, we cannot efficiently compute the three-point function of loci, and thus neglect its contribution to the effective additive weights. This is equivalent to assuming that the three-point correlations of the loci are subleading to the two-point correlations.

***S1.2.2. Tensor inputs to SLED.*** SLED operates most naturally on GP data, taking in a matrix of genotypes and outputting a matrix of phenotypes, as described in the main text. However, as shown in SI S1.2.1, SLED also defines a set of "global" additive and epistatic weights obtained by marginalizing over the latent modules. Thus, in the loss function, one can replace the MSE between the predicted and observed phenotypes with the reconstruction error between a pre-fit set and the effective set of additive and epistatic weights. These ground-truth weights could be inferred by a more traditional statistical process; for example, the forward search algorithm described in SI S2.2 (1). In this setting, SLED effectively acts as a tensor decomposition algorithm, finding a low-dimensional and nonlinear factorization of the effective weights in terms of a small number of modules.

**S1.3. Synthetic data.** Here we discuss the tests on synthetic data in more detail. The goals of these tests were to 1) confirm that SLED could accurately recover sparse, modular structure from GP data designed to match the model's prior; 2) explore the outputs of SLED on data explicitly designed not to match the model's prior; 3) separate the effects of sparsity and modularity; and 4) test the stability of the model across training runs with different random initializations.

As discussed in the main text, we generated synthetic phenotype functions by drawing weights from different generative processes. The weights were 1) sparse and modular, 2) modular but non-sparse (dense), 3) sparse but non-modular, and 4) weighted sums of fifth-order interactions between loci, as a null baseline. All datasets had $N = 100,000$ individuals, $L = 100$ random loci encoded in the $\pm1$ basis, and $E = 20$ environments. We added Gaussian noise with 10% the variance of the phenotypes to each dataset and z-scored the phenotypes and mean-centered the genotypes, as discussed in SI S1.1.2.

When present, sparsity was set such that $W^{(1)}$ had approximately 25% nonzero entries and $W^{(2)}$, which was a smaller matrix, had approximately 50% nonzero entries. The densities of $W^{(1e)}$ and $W^{(2e)}$ were scaled down by $1/L$ ($L$ the number of loci) and $1/M$ ($M$ the number of modules), respectively, relative to the additive weights to make the number of nonzero weights comparable between additive and epistatic effects. Epistatic weights were drawn to match the linked-epistasis constraint, i.e. the probability that an interaction between loci $j$ and $k$ in module $n$ was nonzero was given by

$$P(\left|W^{(1e)}_{njk}\right| > 0) = \exp\left[-\beta_1/\left|W^{(1)}_{nj}W^{(1)}_{nk}\right|\right] \quad \text{[S14]}$$

with $\beta_1$ set to enforce the desired sparsity of the epistatic tensor. An analogous distribution was used for the second layer weights:

$$P(\left|W^{(2e)}_{enm}\right| > 0) = \exp\left[-\beta_2/\left|W^{(2)}_{en}W^{(2)}_{em}\right|\right]. \quad \text{[S15]}$$

All weights, when nonzero, were drawn from $\mathcal{N}(1, 0.5)$. We then applied the gauge-fixing procedure described in SI S1.1.6 to the synthetic weights.

The modular datasets 1) and 2) had $M = 10$ modules. The sparse, non-modular datasets had the same sparsity as described above, but had $M = L = 100$ modules.

Finally, in the fifth-order datasets, each phenotype was a random weighted sum of 500 random interactions, each of five random loci.

Next, we fit the models to the synthetic data and selected the sparsity parameters via the grid search procedure described in SI S1.1.5. As discussed in the main text, we then computed the models' predictive accuracy and ability to recover the ground-truth weights, averaged across five distinct datasets of each type. For one choice of each type of dataset, we also compared 10 models across replicate fits to that dataset by computing the pairwise cosine between models for each set of weights, after gauge-fixing and aligning the modules using a Hungarian matching algorithm (2).

The performance of SLED across these metrics for different synthetic datasets is discussed in the main text. Briefly, as expected, SLED could reliably and accurately recover sparse, modular structure in data drawn from the model's prior. SLED could predict non-sparse, modular datasets well, but struggled to recover the ground-truth weights. SLED struggled to predict and recover sparse, non-modular data, intuitively because of the forced compression through a small number of modules. SLED could not predict the fifth-order phenotypes, as expected; this null baseline provided a sanity check. Replicate fits across random seeds to data not matching the model's prior were much less consistent than fits to the sparse, modular data, providing a way to test for sparse, modular structure in empirical data, where the ground-truth weights are unknown.

**S1.4. Yeast data.**

***S1.4.1. Genome compression.*** The yeast data (1) consist of the relative growth rates of $\sim 100\,000$ F1 segregants from a cross between the BY and RM yeast strains. The segregants differ at $\sim 42,000$ single nucleotide polymorphisms (SNPs). As discussed in the main text, linkage disequilibrium makes most of the loci in the genome redundant. Therefore, we are able to compress the genotypes via the following greedy algorithm. Starting with the first locus, we compute the Pearson correlation across individuals between that locus and each subsequent locus. Proceeding along the genome, we retain the first locus whose absolute correlation with the first locus falls below a fixed cutoff (0.94); this locus then becomes the reference against which the following loci are compared, and the procedure repeats until the end of the genome. This compression reduced our input from 42,000 to 1,429 loci. We also removed loci linked to constructs involved in the experimental protocol. In Fig. S2, we show the selected loci.

Because the genotypes are produced via a cross between two yeast strains, we encode the loci in the $\pm1$ basis; i.e., there is no preferred or "wild-type" allele. For other datasets where a wild-type does exist, e.g. double deletion collections, the 0/1 basis might be more appropriate.

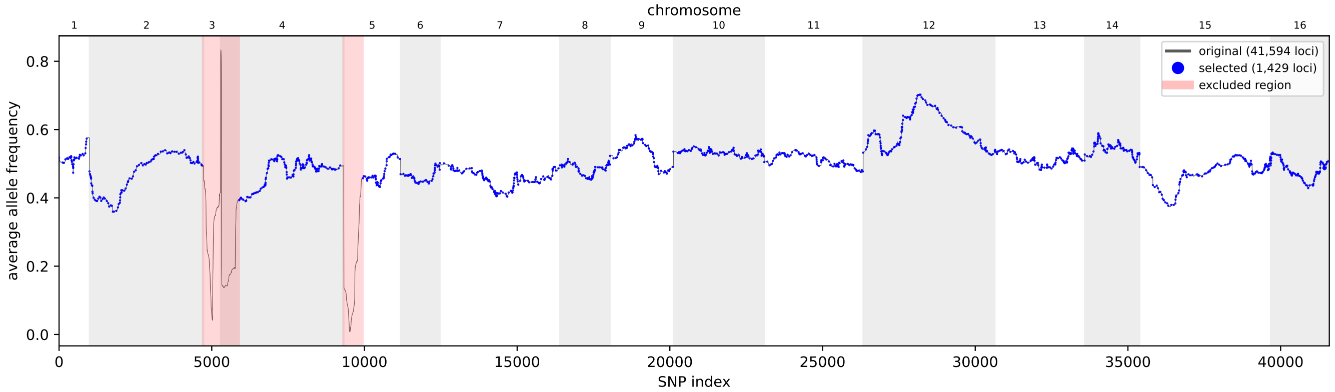


**Fig. S2.** Genome compression for SLED. Plotted is the average allele frequency versus SNP index. The full genotypes consisting of $\sim$ 42,000 loci are compressed to 1,429 loci following the procedure described in the text. The blue points show the selected loci. Two regions containing experimental constructs, highlighted in red, are excluded. The first excluded region includes all of chromosome 3.

***S1.4.2. Train, validation, and test split.*** The training set consists of 85,000 genotypes, of which 90% (76,500) are used for training and 10% (8,500) for validation and early stopping. The remaining 10,235 genotypes are the test set, not seen during training.

***S1.4.3. Convergence with number of training samples.*** There is a practical need to understand how many samples are required to uncover modular structure in the GP map. Here, we empirically explore this question by training models with different numbers of samples $N$. In Fig. S3, we plot versus the size of the training set, averaged over five seed-matched replicates, the mean-squared error of the models as well as the alignment to the full models trained on $N = 85,000$ segregants. Good convergence is seen by $N \approx 30,000$ training samples.

***S1.4.4. Stability of replicate fits to yeast data.*** Training SLED is stochastic, due both to its random initialization and the dynamics of stochastic gradient descent. Nonetheless, SLED learns a highly reproducible structure across replicate fits. In Fig. S4, we show four representative fits, all trained with the same hyperparameters. It is instructive to focus on the top left panel of each model, the additive module activations. As can be seen, the first five modules are quite reproducible. Occasionally, the model will trade higher MSE for a lower regularization cost, combining some of the smaller modules into three or four larger modules. Similarly, there is some variability in resource allocation for smaller modules, e.g. whether or not to learn a separate module for caffeine on the YNB base. Despite this variability, the MSE across 50 replicate models varies by only $\sigma \sim \pm 3.5\%$; gene annotations also do not vary substantially between models.

The visual similarity of the models can be quantified by the same bootstrap procedure discussed in the synthetic data sections. In Fig. S5, we show the heatmaps of the pairwise cosine similarity for each of the four sets of weights across replicate fits. The models are sorted in order of increasing test MSE. This clearly shows that the models are organized into a few distinct subfamilies; the variation between families reflects differing resource allocation to the small modules. The particularly high average similarity of the $W^{(2)}$ weights is indicative of reproducible modular structure. Note that, to account for linkage, weights in the first layer are first smoothed with a Gaussian kernel before the similarity metrics are computed.

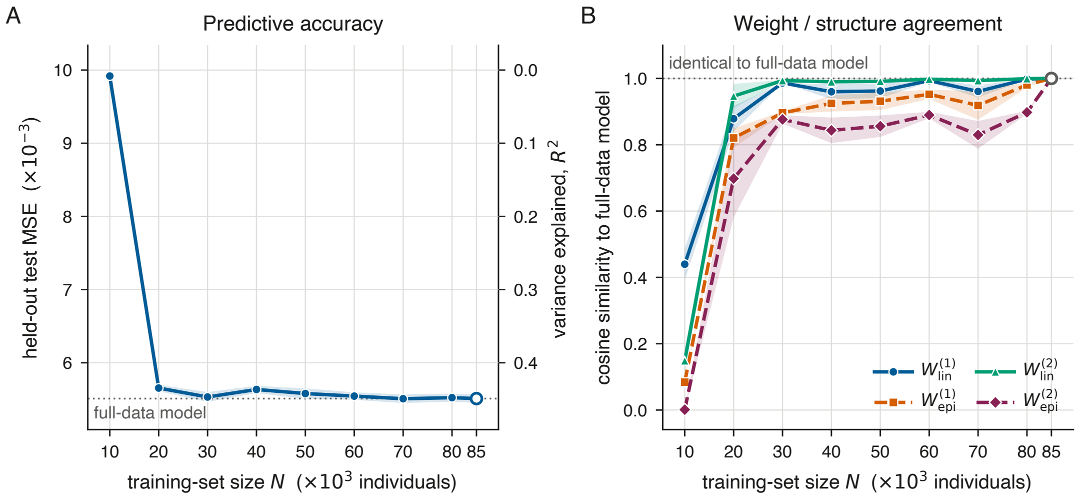


**Fig. S3.** Model convergence with number of samples $N$. **(a)** Mean test MSE versus training set size, shading indicates SEM. **(b)** For each set of weights, mean cosine similarity to the weights learned by the full models trained on $N = 85,000$ segregants, shading indicates SEM. The first layer weights (genes to modules, both additive and epistatic) are smoothed by a Gaussian kernel to account for linkage between nearby loci.

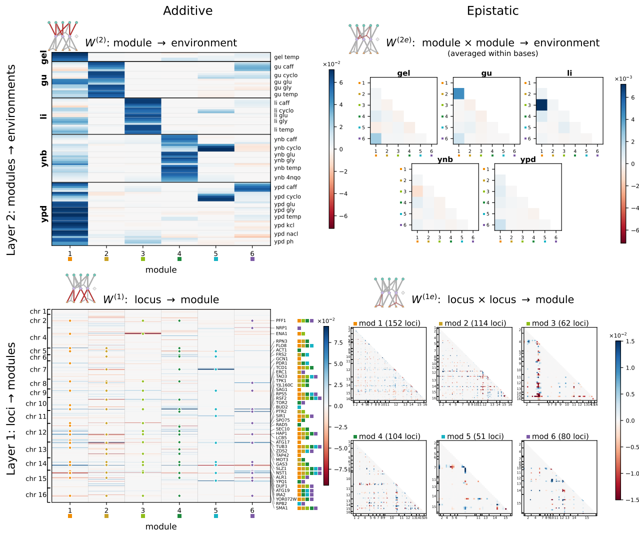


**(a)** Seed 7.

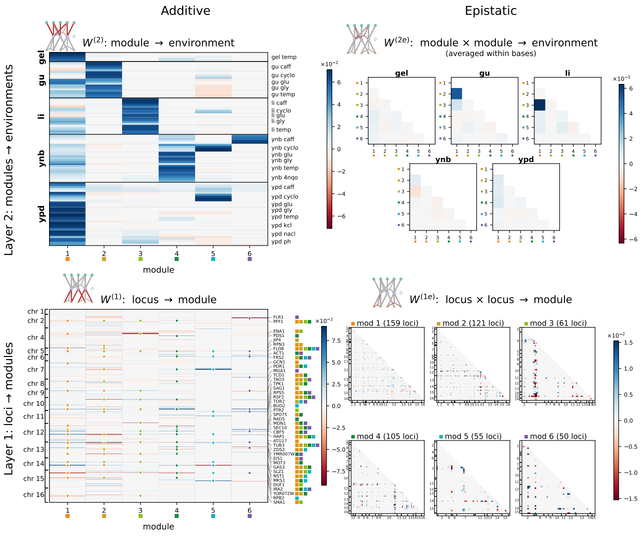


**(b)** Seed 41.

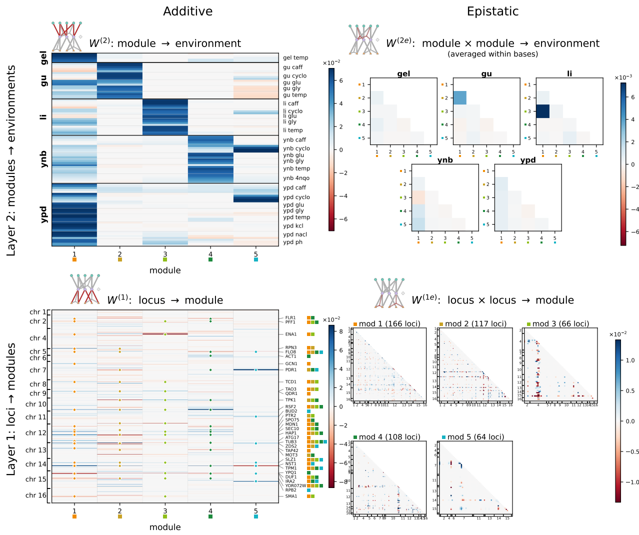


**(c)** Seed 19.

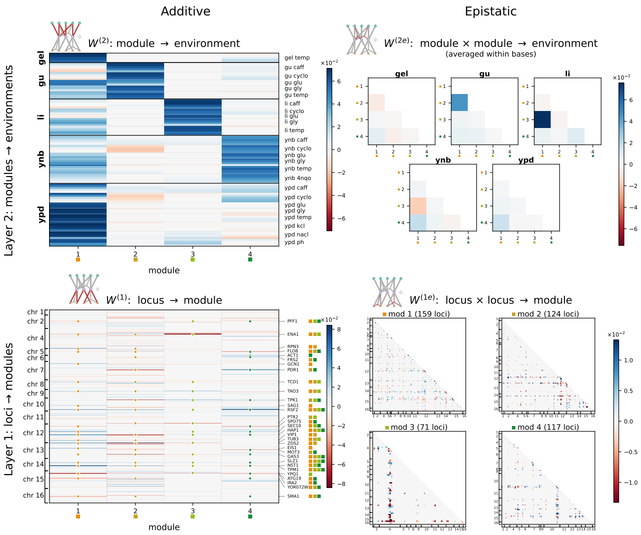


**(d)** Seed 32.

**Fig. S4.** Representative replicate SLED fits to the yeast data, starting from different random initializations. The MSE of the models increases from panels (a) to (d) as the number of learned modules decreases. Seeds correspond to the ones shown in Fig. S5.

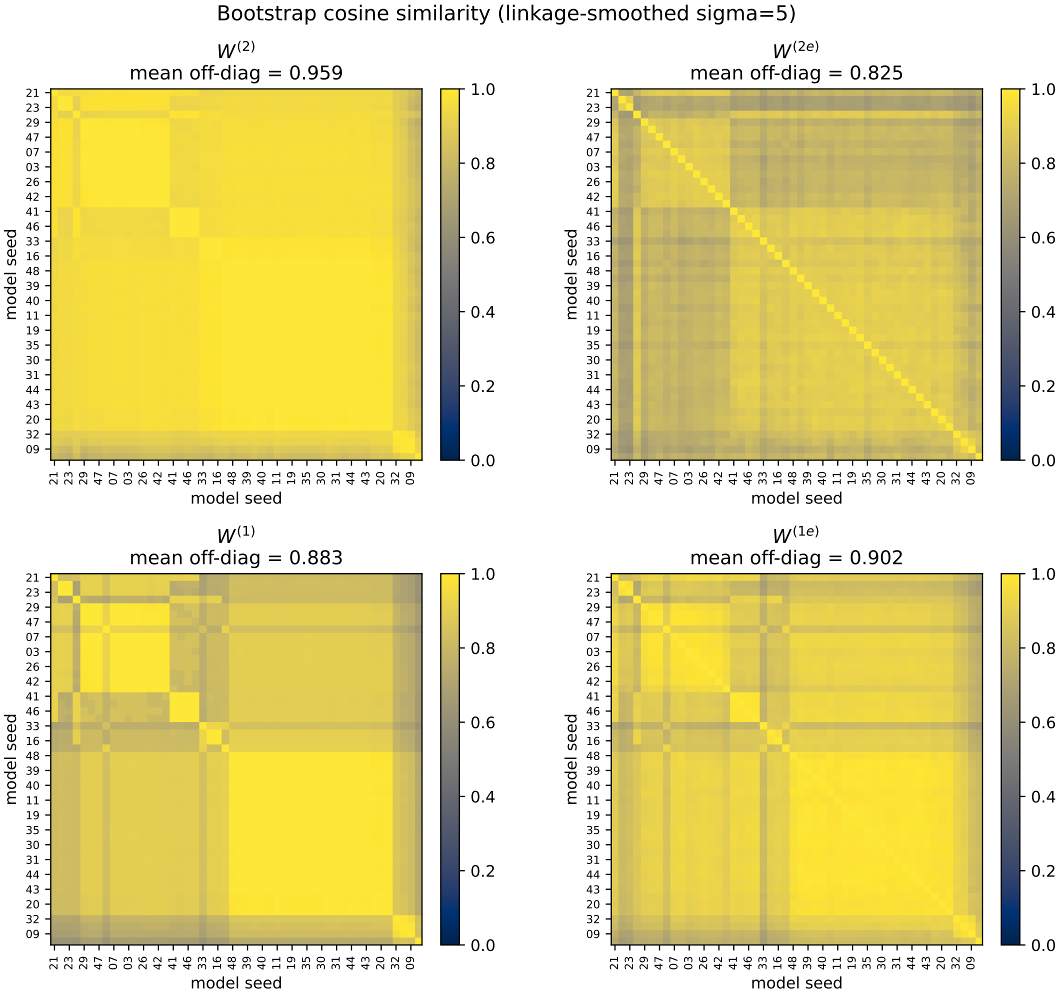


**Fig. S5.** Cosine similarity of replicate fits of SLED to the yeast data. For each of the four sets of weights, the pairwise cosine similarity between the weights learned by each of 50 models is shown. The models are sorted in order of increasing test MSE. The first layer weights (genes to modules, both additive and epistatic) are smoothed by a Gaussian kernel to account for linkage between nearby loci.

#### S2. Supplementary text for GTD

**S2.1. Loss function derivation.** To derive the loss function for our model, we compute the maximum *a posteriori* (MAP) estimate under our prior assumptions about the learned graphs and weights. For one of $E$ total environments, we assume that there are $r$ components such that

$$T^e = \sum_{k=1}^{r} \beta_k^e G_k + \varepsilon^e, \text{ where } \varepsilon^e \sim \mathcal{N}(0, \sigma_N^2 I). \quad \text{[S16]}$$

Using Bayes' Rule, and assuming independence of graph and weight priors, we compute the posterior

$$\begin{aligned} p(G, \beta \mid T) &= \frac{p(T \mid G, \beta) p(\beta, G)}{p(T)} \\ &\propto p(T \mid G, \beta) p(\beta) p(G). \end{aligned}$$

For each environment, $T^e$ is a symmetric matrix with zero diagonal. Since $G$ graphs are also of this form, Eq. (S16) can be equivalently expressed using the flattened upper triangles of the $T^e$ and $G_k$ matrices, which give vectors of length $m := \binom{L}{2}$. Letting $T^e$ and $G_k$ represent these vectors, the likelihood is

$$\begin{aligned} p(T^e \mid G, \beta^e) &= (2\pi\sigma_N^2)^{-\frac{m}{2}} \exp\left(-\frac{1}{2}\left(T^e - \sum_k \beta_k^e G_k\right)^T \frac{1}{\sigma_N^2} I \left(T^e - \sum_k \beta_k^e G_k\right)\right) \\ &= (2\pi\sigma_N^2)^{-\frac{m}{2}} \exp\left(-\frac{1}{2\sigma_N^2} \|T^e - \sum_k \beta_k^e G_k\|_2^2\right), \end{aligned}$$

which we combine over (assumed independent) environments as

$$p(T \mid G, \beta) = \prod_e p(T^e \mid G, \beta^e) = (2\pi\sigma_N^2)^{-\frac{mE}{2}} \exp\left(\frac{-1}{2\sigma_N^2} \sum_e \|T^e - \sum_k \beta_k^e G_k\|_2^2\right).$$

Taking the negative log of both sides gives the negative log likelihood

$$-\log(p(T \mid G, \beta)) = \frac{1}{2\sigma_N^2} \sum_e \|T^e - \sum_k \beta_k^e G_k\|_2^2 + C,$$

where $C = \frac{mE}{2} \log\left(2\pi\sigma_N^2\right)$ is a constant that does not depend on $G$s or $\beta$s. This yields the sum of squared errors (SSE) term of our loss function.

Now, we consider the priors on $G$s and $\beta$s. For the $\beta$ weights, we use a Laplace prior to encourage sparsity across graphs—that is, we expect only a small number of graphs $G_k$ to be active in any given environment. For a single weight $\beta_k^e$,

$$\beta_k^e \sim \text{Laplace}(0, b) \quad \text{implies that} \quad p(\beta_k^e) = \frac{1}{2b} \exp\left(-\frac{|\beta_k^e|}{b}\right).$$

Assuming independence across $k$ and $e$,

$$-\log p(\beta) = \sum_e \sum_k \frac{|\beta_k^e|}{b} + C_\beta = \lambda_\beta \sum_e \sum_k |\beta_k^e| + C_\beta,$$

where $\lambda_\beta = \frac{1}{b}$ controls the sparsity of graph activations across environments, and $C_\beta = rE \log(2b)$.

For a graph $G_k$, let $V_k$ denote its vertex set and $E_k$ denote its edge set. We assume that each vertex is chosen with probability $p$. It follows that

$$\begin{aligned} p(V_k) &= p^{|V_k|}(1-p)^{L-|V_k|}, \\ -\log(p(V_k)) &= |V_k| \log\left(\frac{1-p}{p}\right) + C_V, \end{aligned}$$

for $C_V = -L\log(1-p)$. We define $\lambda_V := \log\left(\frac{1-p}{p}\right)$. This parameter controls the size of the learned interaction modules, with higher values favoring smaller subsets of participating loci.

Turning to our prior on the edges within a fixed vertex set of $G_k$, we assume that each edge is chosen with a probability of $q$, and that when an edge does exist, it can take on any real value with equal likelihood. To ensure model identifiability, however, we require the edge weights for each graph to be normalized. These assumptions yield a spike-and-slab prior on edges.

To derive such an edge prior, we define an indicator $Z_k$ for $G_k$, where

$$(Z_k)_{ij} = \begin{cases} 1 & \text{if } (G_k)_{ij} \neq 0 \\ 0 & \text{else.} \end{cases}$$

It follows that

$$p((Z_k)_{ij} \mid V_k) = q^{(Z_k)_{ij}}(1-q)^{(1-(Z_k)_{ij})},$$

so, given the vertex set $V_k$ of $G_k$,

$$\begin{aligned} p(Z_k \mid V_k) &= \prod_{i<j; i,j \in V_k} q^{Z_{k,ij}}(1-q)^{(1-Z_{k,ij})} \\ &= q^{|E_k|}(1-q)^{\left[\binom{|V_k|}{2} - |E_k|\right]}. \end{aligned}$$

Thus,

$$-\log(p(Z_k \mid V_k)) = |E_k| \log\left(\frac{1-q}{q}\right) + \binom{|V_k|}{2} \log\left(\frac{1}{1-q}\right).$$

Even among participating loci in a module, we may expect a smaller number of those interactions to be active. We define $\lambda_E := \log\left(\frac{1-q}{q}\right)$. This controls the sparsity graphs have within their vertex sets. Finally, we define $\lambda_{V2} := -\log(1-q)$. This term arises from accounting for the number of possible edges a graph can have with respect to its vertex set—since the term is a quadratic penalty on vertices, it favors many small, modular graphs over one large graph.

Combining the vertex and edge priors, assuming all graphs are independent and that edge weights are uniform on the unit $\ell^2$ sphere, we have

$$\begin{aligned} p(G) = \prod_k p(G_k) &= \prod_k p(V_k)\, p(Z_k \mid V_k)\, p(\text{edge weights} \mid Z_k, V_k) \\ &= C_G \prod_k p(V_k)\, p(Z_k \mid V_k), \end{aligned}$$

where $C_G$ is taken as a constant for our modeling assumptions. Hence,

$$-\log p(G) = \sum_k \left[\lambda_V |V_k| + C_V + \lambda_E |E_k| + \lambda_{V2} \binom{|V_k|}{2}\right] - \log(C_G).$$

Combining the priors with the likelihood, and ignoring constants, we obtain the following loss function:

$$\mathcal{L}(T; G, \beta) = \frac{1}{2\sigma_N^2} \sum_e \|T^e - \sum_k \beta_k^e G_k\|_2^2 + \sum_k \left[\lambda_V |V_k| + \lambda_E |E_k| + \lambda_{V2} \binom{|V_k|}{2}\right] + \lambda_\beta \sum_e \sum_k |\beta_k^e|.$$

As written, each term in the loss depends differently on the number of environments, loci, and learned graphs. So, we rewrite our loss function to make the terms within it comparable, and have the same hyperparameter scale across datasets. To do so, we define $\alpha := \frac{2r\sigma_N^2}{Em}$ and let $\tilde{\lambda}_\beta = \alpha E \lambda_\beta$. Writing $\|\beta\|_1 := \sum_e \sum_k |\beta_k^e|$, the loss function can be rewritten to preserve the minimizer as

$$\tilde{\mathcal{L}}(T; G, \beta) = \frac{1}{Em} \sum_e \|T^e - \sum_k \beta_k^e G_k\|_2^2 + \alpha \left(\frac{1}{r} \sum_k \left[\lambda_V |V_k| + \lambda_E |E_k| + \lambda_{V2} \binom{|V_k|}{2}\right]\right) + \tilde{\lambda}_\beta \left(\frac{1}{Er} \|\beta\|_1\right).$$

Since $|V_k|$ and $|E_k|$ are hard counts, they are not differentiable as written. So, we use sigmoid gating as a soft approximation of these counts when implementing the loss function (see SI S2.3.2).

We now provide some intuition for the behavior of these parameters. A value of $p < 0.5$ yields $\lambda_V > 0$, which penalizes the addition of vertices, whereas $p > 0.5$ gives $\lambda_V < 0$, rewarding additional vertices. The same intuition applies to $q$ with respect to edge penalization. To understand the $\binom{|V_k|}{2}$ term, we consider the relationship between vertex and edge sets. The corresponding $\lambda_{V2}$ parameter increases monotonically with $q$, penalizing a term proportional to $|V_k|^2$. As such, increasing edge density via $q$ favors smaller vertex sets. This encourages well-connected graphs that are dense within a small vertex set, rather than large, diffuse graphs. For fixed $p$ and $q$, the strength of $\alpha$ dictates the size of the graph penalty term relative to the MSE and $\beta$ $\ell_1$ terms. Larger $\alpha$ encourages a stronger penalty on graph structure, and smaller $\alpha$ prioritizes minimizing the reconstruction error. Finally, $\tilde{\lambda}_\beta$ is a tunable parameter that controls the extent to which the $\beta$ weights are sparse.

**S2.2. Generating GTD's input.** The GTD method assumes an input of a tensor of epistatic weights stacked across environments. In cases where these weights are not directly known (e.g. measured from a double deletion experiment), we must infer them from given GP data. To do so, we use an external pre-processing procedure called forward search. Full details of this method can be found in Gopalakrishnan et al. (1), but we briefly describe the algorithm here. Forward search is a greedy algorithm that iteratively adds terms to an additive and pairwise epistatic model, selecting at each step the term that minimizes the summed squared residual across environments. It starts by adding loci according to their additive effects to fit a linear model. Once the additive model is fit, the procedure greedily adds epistatic terms between the selected additive-effect loci. Hence, epistatic weights are only assigned to loci with additive effects. While the locus set is shared across environments, additive and epistatic effect sizes vary. Running this procedure on the experimental data produces a tensor of epistatic interactions on a subset of 308 loci across the 101 environments.

#### S2.3. Training.

***S2.3.1. Initialization.*** We initialize GTD by using SVD as a warm start for $\{G_k\}$ and $\{\beta_k^e\}$. Intuitively, while these initial guesses are not sparse, they are aligned with dominant axes of variation in the data. We format the input tensor $T$ by flattening the upper triangle of each $L \times L$ per-environment slice into a $\binom{L}{2}$−length vector. This results in an equivalent representation of $T$ as an $E \times \binom{L}{2}$ matrix, $T_{\text{flat}}$. Applying SVD to $T_{\text{flat}}$ gives $T_{\text{flat}} = U\Sigma V^T$, where $U$ and $V$ are orthogonal, and $\Sigma$ is a diagonal matrix of the singular values of $T_{\text{flat}}$. In our framework, we think of the graphs as representing interaction structure, while the beta weights carry the activations of these interaction components across environments. As such, we take the unit-norm columns of $V$ to be the vectorized (i.e. upper triangle) versions of the initial graphs $\{G_k\}$, and the columns of $U\Sigma$ to be the $\{\beta_k^e\}$ weights across environments for a given graph.

To determine the number of learned graphs, we compute the rank of $T$ needed to explain 90% of the variance using SVD. This rank $r$ acts as a ceiling for the number of learned graphs; depending on the sparsity regularization strength, GTD may not use all $r$ graphs.

***S2.3.2. Optimization.*** We use gradient descent with an Optax Adam optimizer on the loss function derived in SI S2.1. As written, the loss function is not differentiable, since it uses hard counts for vertices and edges. Therefore, we use sigmoid gating as a soft approximation for these counts.

For edges, or interactions between loci, we replace the hard indicator of the activity of entry $(i, j)$ of graph $G_k$ with a sigmoid gate on its magnitude:

$$g_{ij}^{(e)} = \sigma\big(t\,(|G_{k,ij}| - \tau)\big) = \frac{1}{1 + e^{-t(|G_{k,ij}|-\tau)}}.$$

The center $\tau$ determines the threshold magnitude for whether or not an edge is considered active. The steepness $t$ controls the model's confidence in whether an edge participates, and thus the extent to which edge assignment can change during optimization. Summing $g_{ij}^{(e)}$ over the upper triangle of $G_k$ gives a soft edge count $\widehat{|E_k|}$, replacing the hard count $|E_k|$ in the loss function.

For vertices (loci), we first compute a soft degree for locus $i$, i.e. the average edge gate over its $L - 1$ possible interactions:

$$d_i = \frac{1}{L-1} \sum_{j \neq i} g_{ij}^{(e)}.$$

We then apply an analogous sigmoid gate to this soft degree:

$$g_i^{(v)} = \sigma\big(s\,(d_i - \kappa)\big) = \frac{1}{1 + e^{-s(d_i-\kappa)}},$$

where the center $\kappa$ is the fraction of possible edges a locus must softly participate in to be considered active, and the steepness $s$ plays the same role as $t$ does for edges. Summing $g_i^{(v)}$ over all loci gives a soft vertex count $\widehat{|V_k|}$, replacing the hard count $|V_k|$ in the loss function.

While this introduces four additional hyperparameters $(\tau, t, \kappa, s)$, we found that GTD works well across a variety of parameter choices, so we fix the edge and vertex centers at $\tau = \kappa = 0.005$, and the edge and vertex steepnesses at $t = s = 5$ (see SI S2.5 for a full parameter sweep).

***S2.3.3. Pre-processing.*** SLED z-scores phenotypes in each environment, rescaling them to have a fixed variance of 0.01 (see SI S1.1). To match this convention, we compute how z-scoring the phenotypes transforms the epistatic weights that serve as GTD's input. In a given environment $e$, we can write the fitness (phenotype) of genotype $g$ as

$$y^e = C + \sum_i f_i^e g_i + \sum_i \sum_{j<i} f_{ij}^e g_i g_j.$$

For per-environment mean $\mu^e$ and standard deviation $\sigma^e$, z-scoring gives

**Table S1. Hyperparameter grid used in the model selection sweep.**

| Parameter | Values | Count |
|---|---|---|
| $p$ | $0.01,\ 0.05,\ 0.1,\ 0.2,\ 0.5$ | 5 |
| $q$ | $0.01,\ 0.1,\ 0.3,\ 0.6,\ 0.9$ | 5 |
| $\tilde{\lambda}_\beta$ | $1\times10^{-6},\ 3.98\times10^{-6},\ 1.58\times10^{-5},\ 6.31\times10^{-5},\ 2.51\times10^{-4},\ 1\times10^{-3}$ | 6 |
| $\alpha$ | $1\times10^{-7},\ 6.31\times10^{-7},\ 3.98\times10^{-6},\ 2.51\times10^{-5},\ 1.58\times10^{-4},\ 1\times10^{-3}$ | 6 |

$$\frac{y^e-\mu^e}{\sigma^e}=\frac{C-\mu^e}{\sigma^e}+\frac{1}{\sigma^e}\sum_i f_i^e g_i+\frac{1}{\sigma^e}\sum_i\sum_{j<i} f_{ij}^e g_i g_j.$$

As such, the epistatic weights $f_{ij}^e$ are rescaled by $\frac{1}{\sigma^e}$ under z-scoring. To match SLED's target phenotype variance of 0.01, we scale each per-environment slice of $T$ by $\frac{0.1}{\sigma^e}$. This gives each environment equal weight in the loss at the phenotype level, rather than the level of the epistatic weights themselves.

**S2.4. Model selection.** To explore the space of models produced by GTD, we performed a hyperparameter grid sweep of $\{\alpha, p, q, \tilde{\lambda}_\beta\}$. On this dataset, the maximum number of graphs was $r = 15$, which is fixed across this sweep (see SI S2.3 for details on determining rank $r$). The ranges of each parameter are given in Table S1 below, for a total of 900 configurations.

After training each model, we chose a candidate for visualization and further analysis by balancing sparsity in the learned graphs and weights with reconstruction error. We prioritize weight sparsity first, since the $\beta$s determine the activity of a module across environments, while graph sparsity offers secondary refinement of which loci participate in each module. Specifically, we first computed the Pareto front for the 2D axes of MSE versus the average $\beta_k^e$ $\ell_1$ value (an estimate of $\beta$-weight sparsity). We then used the Kneedle algorithm (3) to find the "knee" on the graph, or the configuration that offered the best balance between $\beta$ weight sparsity and reconstruction error (Fig. S6a).

We then fixed the $\tilde{\lambda}_\beta$ value from the knee configuration ($1\times10^{-3}$), and ran the same analysis, this time balancing MSE with average vertex set sizes across graphs (Fig. S6b). Computing the Pareto front and the knee yielded the following configuration of the hyperparameters: $\{\alpha, p, q, \tilde{\lambda}_\beta\} = \{1.58\times10^{-4}, 0.5, 0.01, 1\times10^{-3}\}$. This configuration reflects a strong $\beta$ weight sparsity: the selected model uses only 8 out of 15 possible graphs (Fig. 7).

Several of the selected values lie at the boundary of the grid. For $p$, the boundary is a principled choice: when $p > 0.5$, $\lambda_V = \log\left(\frac{1-p}{p}\right) < 0$, so the prior rewards the inclusion of vertices. At $p = 0.5$, the per-vertex penalty vanishes, but graph size remains penalized through $q$. Specifically, when $q = 0.01$, each pair of participating vertices incurs a penalty of $\lambda_{V2} \approx 0.01$. Although the Pareto choices of $q$ and $\tilde{\lambda}_\beta$ are also at the edges of our hyperparameter grid, the selected configuration lies in the interior of both Pareto fronts (Fig. S6). Hence, the sweep reaches sparser solutions, but at the cost of substantially higher reconstruction error.

**S2.5. Robustness.** Here, we assess the robustness of GTD to both small perturbations in the hyperparameters and model initialization. We also assess the robustness of GTD to changes in the sigmoid hyperparameters, which are fixed during optimization (see SI S2.3.2).

For the hyperparameter perturbations, we fixed the four hyperparameters $\alpha, p, q$ and $\tilde{\lambda}_\beta$, described in Section 2.1 of the main text, at a baseline setting informed by our hyperparameter sweep (see SI S2.4). We then varied each individually by $\pm5\%$ and $\pm10\%$ relative to their baseline values, with the remaining hyperparameters fixed. This yielded 16 configurations for comparison to the unperturbed baseline (Table S2).

For the initialization perturbations, we perturbed the initial values of $\{G_k\}$ and $\{\beta_k^e\}$ obtained via SVD by adding random noise with strength equal to 10% of the standard deviations of the magnitudes of their entries. We did this across 15 random seeds.

Finally, to test the robustness of GTD to changes in the sigmoid parameters, we swept each of the four individually over a fixed grid of four settings (see SI S2.3.2). For our sigmoid centers, we sweep $[0.005, 0.05, 0.1, 0.5]$, which explores a wide range of "active" edges and vertices. The sigmoid steepness $t$ (renamed $s$ for vertices) is swept across $[1, 5, 100, 1000]$ (Fig. S7).

For all tests, we compared perturbed models against the baseline using a ratio of MSE, as well as the average cosine similarity of the graphs and per-environment $\beta$ weights. To align graphs for the similarity assessment, we used the Hungarian matching algorithm (2) after dropping unused modules (see SI S2.6).

The sigmoid parameters, swept over a much larger range than the parameters in the other two tests, vary in the extent to which they impact model error and similarity of learned graphs. While the vertex parameters do not affect GTD's performance, the method is sensitive to changes in the edge sigmoid parameters. Intuitively, we expect the algorithm to be sensitive to the edge sigmoid parameters, since edge selection has downstream effects on the vertex sigmoid (via weighted degree), and the penalties in the loss.

To gain intuition for what happens as $\tau$ increases, suppose that a graph $G$ has 200 active edges of roughly equal magnitude, and $t$ is fixed at 5. Using our graph thresholding heuristic (see SI S2.6), each edge has magnitude $\sim 0.07$. For $\tau = 0.005$, the sigmoid output is $g_{ij}^{(e)} = 0.58$, and for $\tau = 0.5$, this decreases to $g_{ij}^{(e)} \approx 0.1$. As such, an average edge at smaller $\tau$ counts about

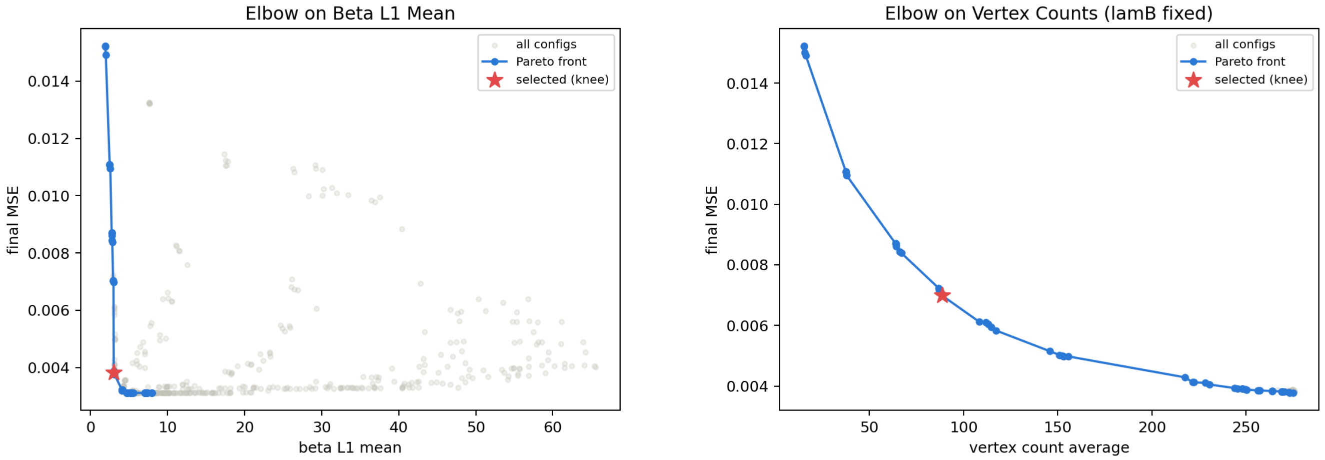


**(a)** **(b)**

**Fig. S6.** Knee-point selection for model choice. (a) Pareto front of MSE versus mean $\beta$-weight $\ell_1$ value, with the selected knee point. (b) Pareto front of MSE versus mean vertex set size, computed after fixing $\tilde{\lambda}_\beta$ at the value selected in (a).

half the time, whereas this likelihood decreases as $\tau$ increases. We conclude that values smaller than 0.5 produce reasonable, robust results.

As for $t$, the instability we see (especially the failure at 100) is reflective of the limits of our optimization. For GTD, we take $t = 5$ as a moderately steep value.

**S2.6. Graph thresholding.** After running GTD, we apply a hard threshold to the edges (interactions) of each graph. This threshold is determined by a heuristic developed to separate noisy, null edges from those carrying meaningful signal. We assume that a graph $G$ is "dead" (i.e. unused in the decomposition) if it has uniform, or near uniform, edge weights; GTD enforces each graph to have unit Frobenius norm, so a uniform edge distribution is just a normalized graph of zeros. Given the size (number of edges) of a graph $n$, we can solve for the magnitude of a uniform edge weight $c$:

$$1 = \|G\|_F^2 = \sum_{i=1}^{n} c^2 = c^2 n,$$

so $c = \frac{1}{\sqrt{n}}$. A locus is considered to be part of a module if it retains at least one surviving edge above this threshold; the procedure therefore reduces both vertex and edge counts. On the yeast dataset we analyze, there are 308 loci, resulting in $\binom{308}{2}$ possible edges. Using our heuristic, the cutoff for a meaningful edge weight is $c \approx 0.0046$. Such a threshold is used in our model analysis in the main text, and in procedures like selecting hyperparameters, where effective vertex and edge set sizes characterize sparsity of a learned model.

After thresholding, if a module has fewer than 10% of the total number of vertices (for the yeast data, this is around 30), we drop those modules from our learned model, which reduces its rank.

**S2.7. Synthetic data.** We designed two complementary sets of synthetic experiments. The first varied graph structure with fixed sparsity (edge density) and graph size (vertex density). This tested the flexibility of GTD across structurally distinct data. The second test varied edge and vertex density with a fixed Erdős–Rényi graph structure to match our loss function (4, 5). This targeted GTD's prior assumptions of sparsity and low-rank structure.

***S2.7.1. Varying graph topologies.*** Our first test suite varied the structure of the underlying generating graphs. For each of 5 graph-generation schemes, we sampled 8 generating graphs with $L = 40$ loci. Each graph had a randomly drawn vertex set (25% of all loci), and edge density of 0.3 when applicable. Edge weights were drawn from a normal distribution, then each graph was rescaled to unit Frobenius norm to match GTD's convention. Per-environment $\beta$ weights were active with probability 0.6, drawn across 50 environments. We combined these synthetic data into a tensor $T$, building each per-environment slice as a linear combination of graphs scaled by $\beta$ weights. Finally, we added symmetric Gaussian noise to $T$, with standard deviation of 10% of that of $T$.

We generated five sets of generating graphs:

- **Erdős–Rényi (ER).** Edges are independently assigned at a fixed probability (4, 5). This structure directly matches GTD's prior.

**Table S2. Robustness of GTD to hyperparameter and initialization perturbations. Values report mean $\pm$ std across all perturbed configurations (hyperparameter perturbation, $n = 16$) or random seeds (initialization perturbation, $n = 15$), relative to the unperturbed anchor model.**

| Metric | Hyperparameter perturbation | Initialization perturbation |
|---|---|---|
| MSE ratio | $0.998 \pm 0.017$ | $1.003 \pm 0.005$ |
| Graph cosine similarity (mean) | $0.999 \pm 0.001$ | $0.995 \pm 0.0004$ |
| $\beta$ cosine similarity (mean) | $0.999 \pm 0.001$ | $0.995 \pm 0.0003$ |

- **Bipartite.** A graph is bipartite if its vertices can be partitioned into two disjoint sets such that every edge joins a vertex in one set to a vertex in the other. This tests whether GTD recovers exact generating graphs via their shared usage across environments, or instead separates structure within a graph, producing a modular solution without exact recovery.
- **Star.** One central "hub" vertex has edges to all other vertices. Recovery of star graphs is important biologically, since hub loci that interact with many other loci are a common epistatic motif.
- **Configuration model.** Vertex degrees are drawn from a power law distribution and then randomly paired (6, 7). This generalizes the star graph structure and tests whether GTD can recover interaction structure beyond a single dominant hub.
- **Mixed.** All four random graph types are combined at once, testing whether GTD recovers each latent graph structure rather than a blurred low-error solution.

GTD recovered the generating graph set across all 5 configurations with a cosine similarity of 1. On the other hand, when we ran SVD, the learned graphs had cosine similarities as low as 0.4 (Figs. 5 and S8). The SVD baseline fared best with the Erdős–Rényi setting, but struggled with star and configuration model graphs.

***S2.7.2. Varying sparsity and disjointness.*** In our second round of synthetic tests for GTD, we focused on varying the sparsity of both the vertex and edge sets. For all of these tests, we used Erdős–Rényi random graphs. We sampled 10 generating graphs with $L = 300$. Per-environment weights were active with probability 0.6, drawn across 100 environments. Analogous to the first test suite, graphs were rescaled to unit Frobenius norm, and we added 10% symmetric Gaussian noise to $T$.

The first axis, edge density, dictates the number of edges a graph has on its support. Before drawing edges, we generated a vertex set according to vertex set sparsity. Once this set was fixed, we drew edges between them randomly with probability $q$. For these tests, we chose a binary setting of "sparse" ($q = 0.3$) versus "dense" ($q = 0.7$). The second axis, vertex density, controls the size of the vertex set. A sparse vertex set uses 10% of the total available vertices (i.e. 30 of 300), whereas a dense one uses 90% of the total available vertices.

We fixed the generating graphs to share the same support, and overlap with each other. While it is possible to generate a tensor from graphs that are largely disjoint, provided that they have small enough (i.e. sparse) vertex sets, such structure is easily recovered by GTD, and is too simple to reflect real biological data. As such, depending on the size of the vertex sets of the generating graphs, we limited the pool of loci that each graph can sample vertices from to force substantial overlap.

There are four combinations of the two axes of sparsity. GTD performed well on both sparse and dense edge configurations. However, GTD struggled with dense, overlapping vertex sets, regardless of edge density (Fig. 5). This suggests that GTD is not designed to handle graphs that are truly not sparse in participation, and that overlap significantly, which aligns with our assumptions that GTD should recover sparse, low-rank structure.

We remark that for both the graph structure and sparsity synthetic testing, we used the same regularization within each test suite. For the first set of tests, we used $\alpha = 1.5 \times 10^{-4}, p = 0.3, q = 0.6, \tilde{\lambda}_\beta = 2.5 \times 10^{-4}$. For the second, we set $\alpha = 1 \times 10^{-5}, p = 0.3, q = 0.2$, and $\tilde{\lambda}_\beta = 1 \times 10^{-3}$. An informal sweep of the space for the dense vertex set configurations (where GTD does not recover the generating graphs) suggested that GTD cannot find the underlying structure regardless of model settings.

#### S2.8. MCPCA comparison.

***S2.8.1. Synthetic comparison to MCPCA.*** In addition to comparing the output of GTD with SVD, we compare GTD to another tensor decomposition method: Multi-context principal component analysis (MCPCA) (8). MCPCA writes $T = \sum_{k=1}^{r} a_k \otimes a_k \otimes b_k$. Here, $a_k \otimes a_k$ (or $a_k a_k^T$) is an $L \times L$ rank-one component, which is one factor, and $b_k$ is an $E$-length vector that encodes how much the factor is used in each environment. For a fixed rank $r$, MCPCA and SVD of the flattened tensor capture the same subspace of $L \times L$ components, but differ in their choice of basis: MCPCA selects rank-one symmetric components $a_k a_k^\top$, whereas SVD returns an orthogonal basis whose elements are linear combinations of these components. Hence, when the underlying components are close to rank-one, MCPCA can recover them individually, while SVD mixes them unless they happen to be orthogonal.

Below, we compare the results of GTD and MCPCA on three synthetic graph topologies: Erdős–Rényi, star, and rank-one graphs (Fig. S9). We note that since MCPCA assumes that the $b_k$ vectors are non-negative, we use the related algorithm, the Multi-Subspace Power Method (MSPM), which is the same procedure, but lifts the non-negativity constraint (9).

Since our synthetic tensors were each generated with 8 graphs (see Section 2.2), we set the number of components for MCPCA to learn to 8 as well. MCPCA is unable to recover the ER graph structure, since the random generating procedure

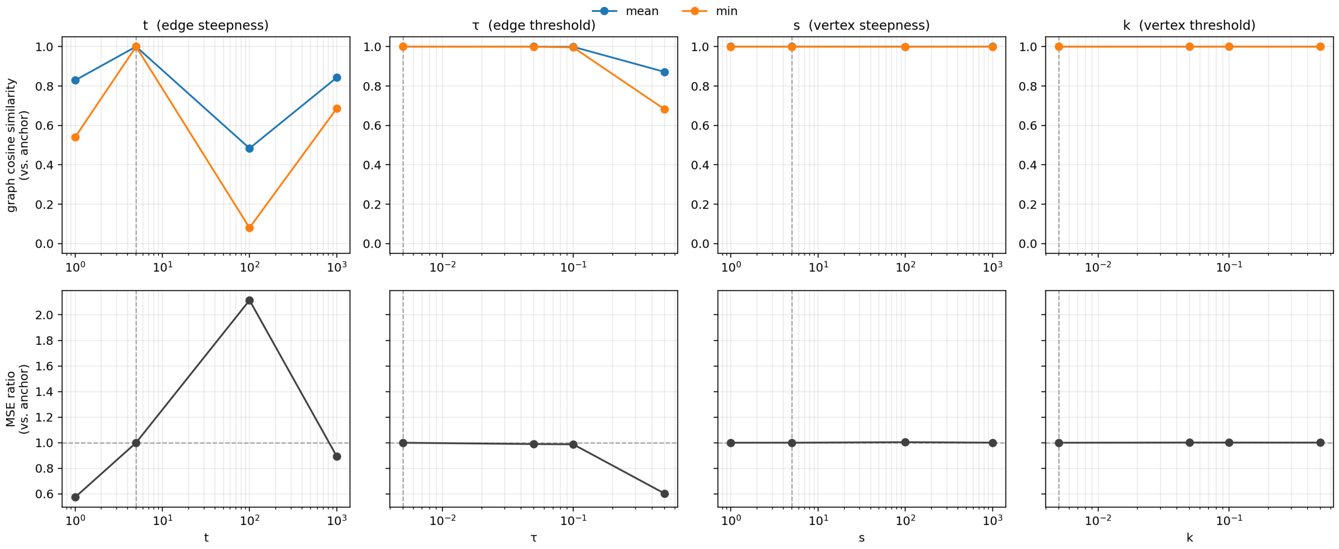


**Fig. S7.** Robustness of GTD to variation in the four model sigmoid parameters. The top row shows cosine similarity of learned graphs from a perturbed configuration to an anchor configuration (the fixed GTD settings), and the bottom row shows the ratio in MSE between the perturbed configuration and the anchor. Each column represents the one-at-a-time sweep of one parameter, holding the others fixed.

often yields high-rank graphs. Hence, to learn the full structure under MCPCA's rank-one constraint would require learning more components than the number of generating graphs.

On the star configuration, MCPCA is much better at recovering the underlying structure, as star graphs are much closer to rank-one. Finally, MCPCA nearly perfectly recovers structure in the rank-one setting, since it directly aligns with the assumption of the method.

On each of the three configurations, GTD performs similarly well, and recovers the generating graph set. The flexibility of GTD to recover various underlying graph structures is due to its prior, which enforces sparsity, but does not have any constraint on structure.

MCPCA takes advantage of tensor structure; it is able to explain directions of variance across subsets of environments. As such, it provides a useful initial pass on the yeast data (see SI S2.8.2). However, it differs from the goals of GTD in that it learns graphs that are rank-one. In our context, this assumption means that in a module, each participating locus must interact with all others. GTD assumes that within a module, structure is more arbitrary: loci may or may not interact.

***S2.8.2. MCPCA on yeast data.*** We also present the results of running the MCPCA method on the yeast dataset, using the same inferred epistatic weight tensor that GTD takes as input (Fig. S10). For stability of the method, we fill the diagonal entries with the absolute sum of their corresponding rows. We run MCPCA using 15 components, which was an arbitrary choice in the ballpark of number of learned modules using SLED, GTD, and SSD; due to the high rank of the epistatic weight tensor, the number of MCPCA components required to capture full signal is likely much higher. We also normalize the learned graph components, and absorb scale into the per-environment weights, to match the convention of GTD.

Because MCPCA finds rank-one components, running it on the yeast data yields components that choose one hub locus, and concentrate other interactions around that. As such, it finds loci that contribute to maximum directions of change in epistatic effects across environments. We remark that filling the diagonal results in fully positive per-environment weights. This is likely because the fill makes each slice $T^e$ diagonally dominant, and hence positive semidefinite, which is consistent with a decomposition $\sum_k b_k^e a_k a_k^\top$ with nonnegative $b_k^e$. Running MCPCA without this fill yields different results. Out-of-the-box, MCPCA is fundamentally different from the methods presented in this work due to the structure of components it finds. As such, it is possible to consider how MCPCA could be adapted to fit the priors we assume on the data; the rank-one components learned could serve as sets of participating loci, and edges could be refit within that support. MCPCA could also be adapted to enforce sparsity in the module-to-environment layer.

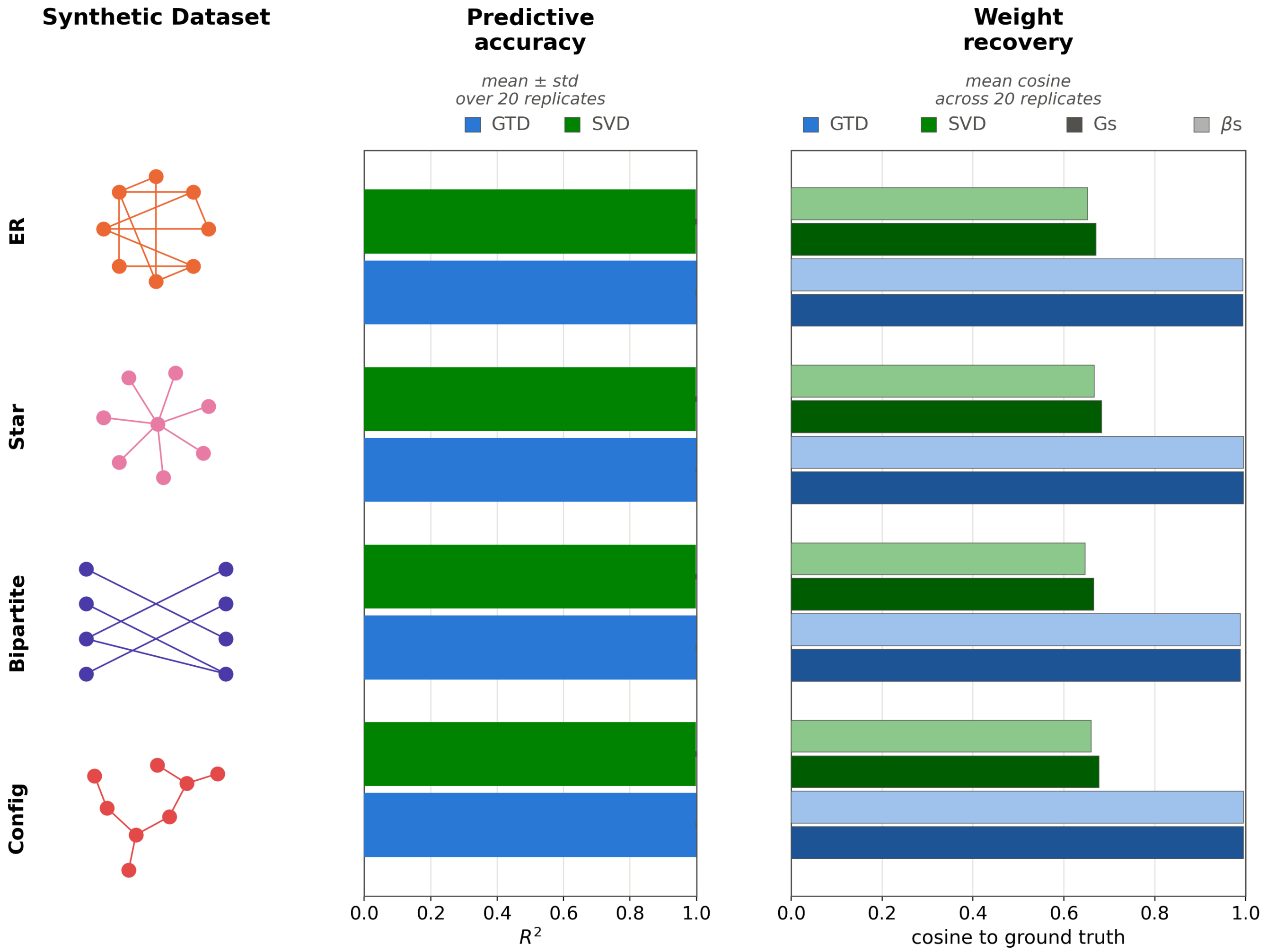


**Fig. S8.** Analysis of the remaining synthetic data configurations not shown in Figure 5. Panel layout and metrics are discussed in Figure 5.

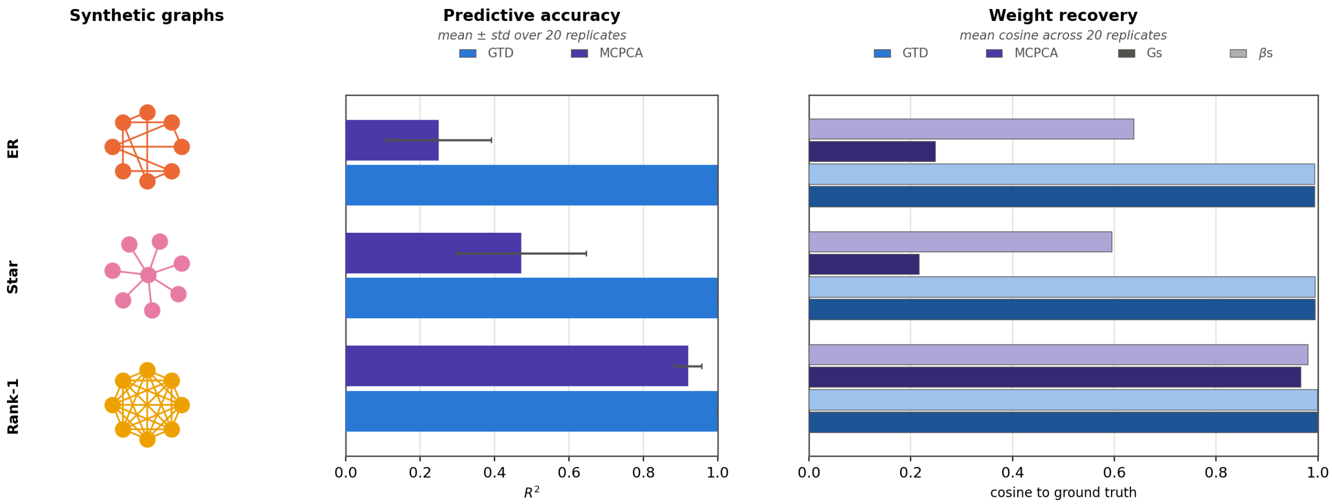


**Fig. S9.** Comparison of GTD and MCPCA on select synthetic data configurations (Erdős–Rényi, star, and rank-one topologies). Panel layout and metrics are discussed in Figure 5.

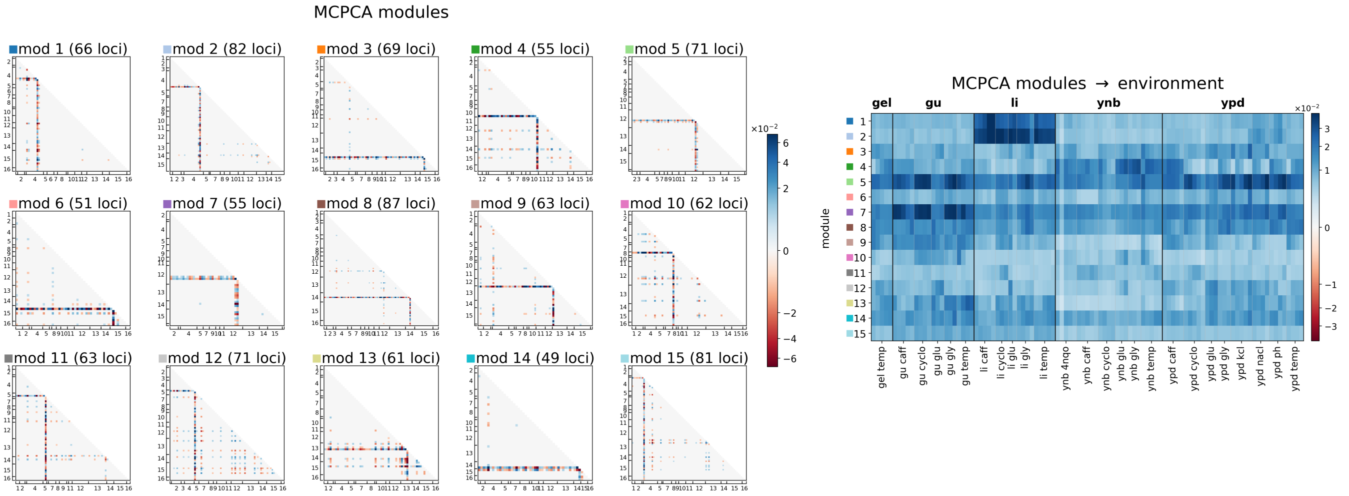


**Fig. S10.** MCPCA applied to the inferred epistatic weights from the yeast dataset. Left: epistatic locus to component map. An interaction is included in a module according to the thresholding procedure used by GTD (see SI S2.6). Loci are labeled by chromosome. Right: module-to-environment map.

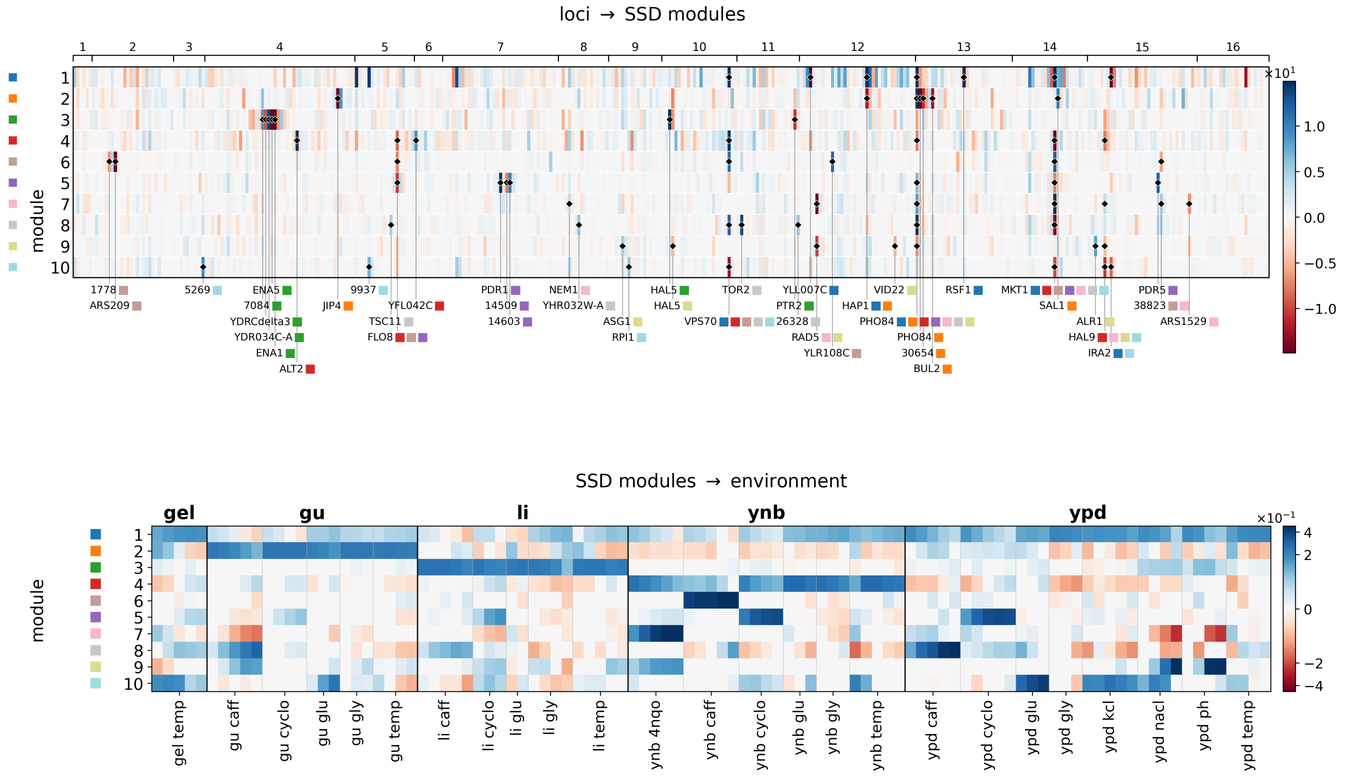


**Fig. S11.** Overview of SSD. SSD applied to the inferred additive effects from the yeast dataset. Top: locus-to-module map. Loci are ordered by genomic position and labeled by chromosome; color indicates each locus's signed weight in each module. The 10 strongest loci in each module are marked and annotated, with colored squares indicating the module(s) they belong to. Bottom: module-to-environment map; color indicates each module's signed weight in each environment. Base media are separated by thick black lines, and perturbations within them by thin gray lines.

#### S3. SSD

Here, we discuss SSD (10), the sparse, low-rank decomposition of additive effects that we compare to SLED and GTD. Unlike SLED and GTD, SSD does not account for or model any epistatic interactions. That is, SSD assumes that loci group into a small set of modules according to their additive effects on phenotypes, rather than how they interact with each other. Further, SSD assumes that fitness effects across environments are linear combinations of these modules, so that modules do not interact with each other. The input to SSD is a pre-inferred matrix of additive fitness effects (see SI S2.2).

By contrast, SLED generalizes SSD by allowing for the nonlinear interactions, both within and between modules. GTD, on the other hand, functions as an extension or epistatic version of SSD, decomposing inferred epistatic fitness effects across environments into a linear combination of sparse interaction graphs.

In Section 3.2, we compare SLED and GTD to SSD. We explore how the learned modular structures depend on what kinds of information a model takes as input. To run SSD on the yeast data, we used the inferred additive weights from the forward search procedure (see SI S2.2); the results are shown in Figure S11. The set of loci found to have significant additive contributions to fitness are a superset of the loci that we input to GTD; in order for forward search to add an epistatic term to its fitness model, the two interacting loci must participate additively. We set the maximum number of modules for SSD at $K = 10$. The method can choose to use all 10, or fewer, depending on the setting of the sparsity hyperparameters. To choose the parameters, we used the procedure outlined in the original SSD paper to search the solution space. This gave one particular setting for the sparsity of the loci to module layer, as well as the module-to-environment layer.
